# Sub-Second Fluctuation between Top-Down and Bottom-Up Modes Distinguishes Diverse Human Brain States

**DOI:** 10.1101/2025.03.12.642768

**Authors:** Youngjai Park, Younghwa Cha, Hyoungkyu Kim, Yukyung Kim, Jae Hyung Woo, Hanbyul Cho, George A. Mashour, Ting Xu, Uncheol Lee, Seok-Jun Hong, Christopher J. Honey, Joon-Young Moon

## Abstract

Information continuously flows between regions of the human brain, exhibiting distinct patterns that dynamically shift across states of consciousness, cognitive modes, and neuropsychiatric conditions. In this study, we introduce Relative Phase Analysis (RPA), a method that leverages phase-lead/lag relationships to reveal the real-time dynamics of dominant directional patterns and their rapid transitions. We demonstrate that the human brain switches on a sub-second timescale between a top-down mode–where anterior regions drive posterior activity–and a bottom-up mode, characterized by reverse directionality. These dynamics are most pronounced during full consciousness and gradually become less distinct as awareness diminishes. Furthermore, we find from simultaneous EEG–fMRI recordings that the top-down mode is expressed when higher-order cognitive networks are more active while the bottom-up mode is expressed when sensory systems are more active. Moreover, comparisons of an attention deficit hyperactivity disorder (ADHD) inattentive cohort with typically developing individuals reveal distinct imbalances in these transition dynamics, highlighting the potential of RPA as a diagnostic biomarker. Complementing our empirical findings, a coupled-oscillator model of the structural brain network recapitulates these emergent patterns, suggesting that such directional modes and transitions may arise naturally from inter-regional neural interactions. Altogether, this study provides a framework for understanding whole-brain dynamics in real-time and identifies sub-second fluctuations in top-down versus bottom-up directionality as a fundamental mechanism underlying human information processing.

## 1 Introduction

Information continuously flows between distinct regions of the human brain [1–12], forming characteristic patterns that dynamically reconfigure across states of consciousness [1, 4, 5], cognitive modes [3, 10, 11], and neuropsychiatric conditions [13–16], and understanding these patterns is essential for unraveling the neural mechanisms underlying the multifaceted nature of human behavior.

A key challenge in neuroscience is to reliably capture and interpret the directionality and flow of information across distributed brain regions. In this context, the conceptual distinction between top-down and bottom-up information flow has emerged as a pivotal framework for understanding inter-regional communication [3, 8–11, 17]. Top-down signals, emanating from higher-order cognitive regions, are thought to modulate sensory processing and guide goal-directed behavior, whereas bottom-up flows, originating in sensory cortices, underpin the encoding of environmental stimuli. Due to limitations in temporal resolution, neuroimaging studies have largely focused on functional networks biased toward these directional modalities, as observed in Blood Oxygen Level-Dependent (BOLD) activity patterns from functional magnetic resonance imaging (fMRI) [3, 17–21]. In contrast, electrophysiological investigations using electroencephalogram (EEG) and electrocorticography (ECoG)—with their superior temporal resolution—have more directly addressed the flow directionality and demonstrated its association with distinct cognitive processes [5, 7–11]. Yet, the dynamics governing the transitions between these directional modes remain largely enigmatic: How do these dominant patterns shift in real time? Moreover, can the transition dynamics observed in brain waves be systematically linked to functional networks derived from BOLD activity?

Traditional measures applied to electrocortical signals—such as traveling wave propagation metrics [8–11] and phase based indices [22, 23]–have provided valuable insights into specific aspects of inter-regional directionality. Numerous phase based indices quantify the probability that the phase of one signal consistently leads or lags another over defined time windows, thereby capturing asymmetric interactions in a time-averaged manner [5, 7, 23]. Similarly, traveling wave analyses, which track the propagation of neural mass activity within selected brain regions, have been linked to distinct cognitive functions, particularly in relation to bottom-up and top-down directionalities [9–11]. Despite their utility, these approaches are constrained by their reliance on time-averaging (in the case of phase-based measures) and their focus on preselected local regions (in traveling wave analyses), limiting our understanding of whole-brain dynamics in real-time.

Here, we introduce a new framework — the Relative Phase Analysis (RPA) — which focuses on the top-down and bottom-up modes and the transitions between them. By exploiting phase-lead/lag relationships in the brain waves, our approach provides a robust method to assess the real-time dynamics across the entire brain, revealing fine-temporal transitions of whole brain directionality patterns. We demonstrate that the human brain continually alternates between two dominant global modes on the scale of approximately 200 ms: a top-down mode, in which frontal regions lead posterior regions in phase, and a bottom-up mode, where posterior regions lead frontal regions. These modes exhibit distinct spatial and temporal profiles across different states of consciousness and cognitive conditions, as revealed by our analyses of diverse electrophysiological and neuroimaging datasets.

First, by applying RPA to EEG data obtained across various levels of consciousness during general anesthesia [24], we identified dominant modes of anterior-to-posterior and posterior-to-anterior directionality between which the brain dynamically transitions. Notably, these characteristic modes and transitions are most pronounced during full consciousness, suggesting that they serve as neural signatures of conscious information processing. As participants lose consciousness, these patterns progressively diminish—yielding a more random profile—a phenomenon that we interpret as indicative of altered information processing.

Second, we examine the functional relevance of the phase dynamics using simultaneous EEG–fMRI data [25]. By correlating these dynamics with task-related BOLD signals, we found that the anterior-to-posterior mode is associated with activity in higher-order cognitive networks, such as the Default Mode Network (DMN) and Fronto-Parietal Network (FPN), whereas the posterior-to-anterior mode correlates with the activity of the Visual Network (VN) and Dorsal Attention Network (DAN). These results lead us to conclude that the anterior-to-posterior mode reflects top-down directionality, while the reverse represents bottom-up directionality, thereby linking EEG dynamic patterns with network-level BOLD activation.

Third, we examined group-level differences in mode transition dynamics using RPA by comparing typically developing individuals with a neuropsychiatric cohort, focusing on the attention deficit hyperactivity disorder (ADHD) inattentive subtype. Our analyses reveal distinct imbalances between reduced front-to-back (top-down) directionality and excessive back-to-front (bottom-up) directionality. This supports the theory that ADHD-inattentive symptoms are associated with a lack of top-down signaling and a failure to inhibit bottom-up processes [26]. Furthermore, the enhanced performance of the RPA-based individual diagnostic predictor compared to the conventional Theta/Beta Power Ratio (TBR) underscores its potential as a more effective EEG biomarker for the ADHD-inattentive subtype using resting-state data. These results emphasize the significance of findings identified through simultaneous EEG-fMRI data, as well as the validity of RPA applications for mental disorders at the cognitive level.

Finally, to further elucidate the mechanisms underlying these dynamic patterns, we simulated relative phase dynamics using a generalized coupled-oscillator model on a structural brain network [27, 28]. The simulations successfully reproduced our empirical findings, demonstrating that distinct directional modes and the transitions between them emerge inherently and spontaneously from interactions among neural masses across the brain network.

Collectively, our study introduces **Relative Phase Analysis** as a comprehensive, real-time framework for understanding whole-brain dynamics, providing new insights into hierarchical processing, brain state transitions, and neuropsychiatric conditions. We reveal that the fluctuating interplay between top-down and bottom-up processing is an intrinsic feature of large-scale brain dynamics, and a fundamental mechanism underlying human information processing. By applying RPA, we uncover levels of consciousness, external stimuli, and pathological conditions influence the delicate balance, paving the way for a deeper exploration of the fundamental mechanisms governing perception, cognition, and consciousness.

## 2 Results

### 2.1 Defining Relative Phase Analysis and comparison with other measures

Throughout the manuscript, we focus on the alpha band (8–12 Hz), as it most clearly exhibits the transition dynamics observed in our study. Similar patterns emerge in other frequency bands—including lower beta (12–20 Hz), theta (4–8 Hz), higher beta (20–30 Hz), delta (1–4 Hz), and gamma (30–50 Hz)—albeit with a progressively reduced prominence of these dynamics (see Supplementary Information Section B for more detail).

We begin by validating the relative phase as a robust measure of directionality in brain activity, by comparing it with amplitude and phase-lag measures, as well as with measures that quantify traveling wave propagation. We will subsequently extend RPA to a real-time, whole-brain framework, thereby overcoming the spatial and temporal limitations of previous approaches.

Brain waves from each EEG channel are decomposed into amplitude and phase components over time using the Hilbert transform, with the phase reflecting the precise timing of a signal within its periodic cycle. We then computed the relative phase for each channel by subtracting the global mean phase (across all channels) at each time point. Next, we weighted the resulting phase by applying the sine function, mapping the values continuously from –1 to +1. In this framework, –1 corresponds to a 90° phase lag, +1 to a 90° phase lead, and 0 is obtained for phase differences of 0° or 180°. This weighting strategy is inspired by phase-based measures that mitigate the heightened sensitivity to noise—arising from factors such as volume conduction—near 0° and 180° differences in phase-lead/lag calculations (Figure 1) [22]. We hypothesize that the phase-lead/lag relationships serve as indicators of the directionality of information flow [5, 8–12, 23].

**Fig. 1.**
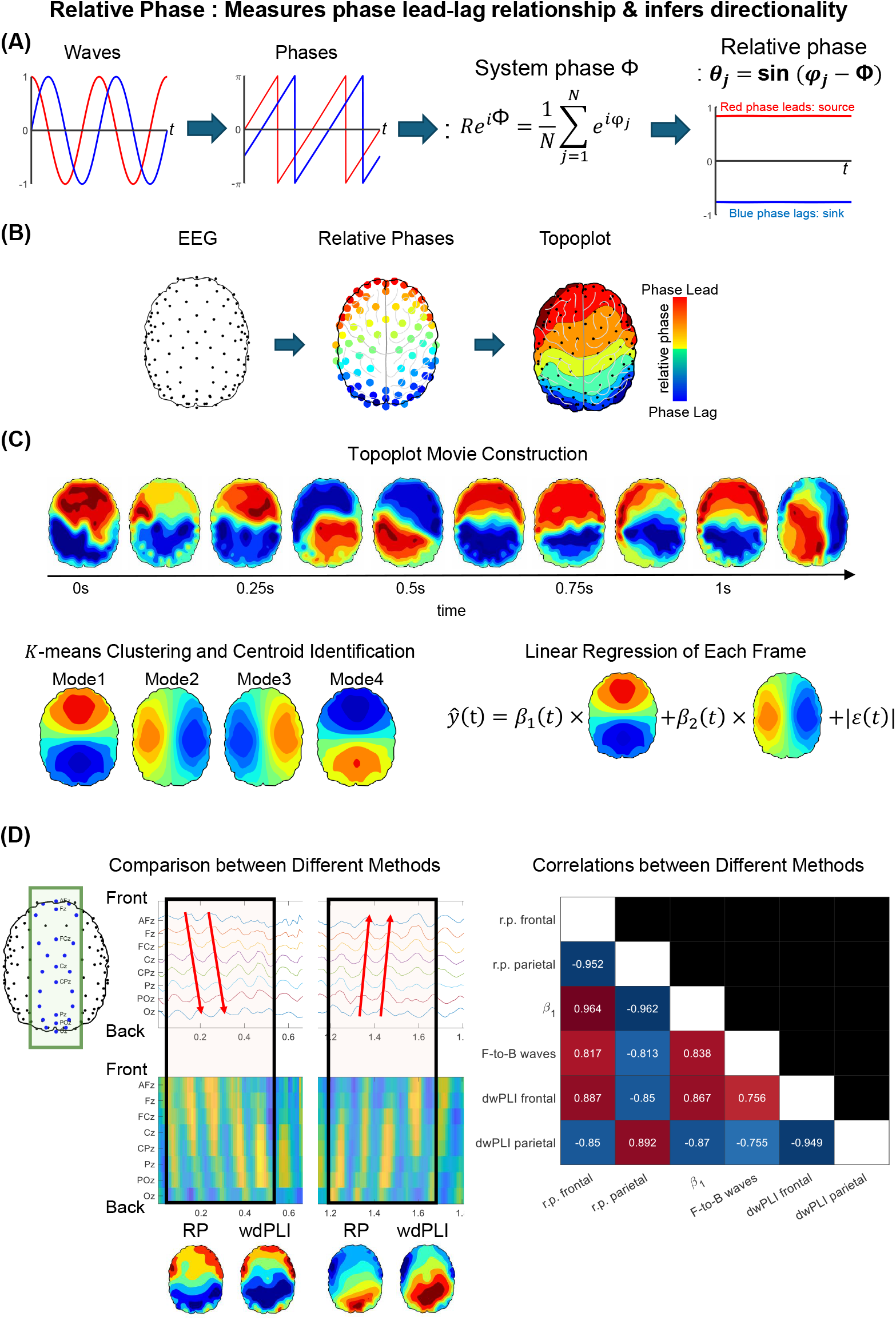
Schematics of relative phase analysis. (A) After performing a Hilbert transform on EEG voltage time series to extract phase information, we calculate the global systems phase by averaging phases of all waves, and subtract them from the phases of each wave. Finally, we take sine of the differences and define the value as the relative phase. (B) After calculating the relative phase from each EEG channel, we spatially extrapolate to construct the topographic map of the relative phase at each time point. (C) By concatenating the topographic map of each time point, we construct the topographic movie of each data set therefore revealing the dynamics of the relative phase. Then, we apply *K*-means clustering across the topographic map of each time point to reveal dominant patterns from the topographic movie. We also apply linear regression taking the centroids of the *K*-means clustering as the vector of regressors, where the topographic map of each time point t becomes the dependent variable ŷ(*t*). (D) We compare the RP (relative phase) measure averaged within time windows against other measures such as wdPLI (weighted directional phase-lag index) and traveling wave measures.

The eyes-closed resting state data from the general anesthesia dataset [24, 29], collected at the University of Michigan (UofM) using 128 electrode channels at a 500 Hz sampling rate, was employed for method comparison. Due to the computational demands imposed by this high-density, high-resolution data, we downsampled the dataset. To ensure that this reduction did not compromise critical information, we determined the optimal time scale for capturing relative phase dynamics (see Figure S7 in the Supplementary Information). We observed that the Shannon entropy of relative phase—computed over time windows of varying sizes—decreases as the window size increases across all frequency bands, indicating that the diversity of relative phase patterns diminishes with broader time averaging (We note that the Shannon entropy of relative phase was computed for each channel first then averaged over all channels). Notably, the alpha band (8–12 Hz) retains its Shannon entropy over longer time scales compared to other bands, up to approximately 100 ms (equivalent to a 10 Hz sampling rate), suggesting that 100 ms is a characteristic time scale for its relative phase patterns. Consequently, in our main analysis, we focus on the alpha wave’s relative phase time series downsampled to 10 Hz to examine brain-state-dependent characteristics. We further note that increasing the downsampling rate to 50 Hz (yielding a time resolution of 20 ms) produces transition dynamics results that are highly consistent (see Supplementary Information Section B.7 for more detail).

The alpha-band filtered data were segmented into 2-second (or 10-second; see Supplementary Information Section B.1 for more detail) time windows, and key metrics were computed. Next, pairwise linear correlations were performed to assess relationships across brain states. Amplitude, derived via the Hilbert transform, was averaged across frontal and parietal regions. Phase-based directionality was quantified by calculating the weighted phase-lag index (wPLI) within each time window, which was then time-averaged and subsequently grouped by frontal and parietal regions. Traveling wave propagation was analyzed using 2D FFT representations and phase gradients computed across selected electrodes traversing the midline of the brain, with the results averaged over time windows. Finally, the relative phase was calculated for all channels and averaged across time windows.

Relative phase showed a strong correlation with measures of information directionality in neural activity (Figure 1D) We found that the relative phase averaged over frontal regions shows a correlation of 0.887 with directional wPLI over frontal regions, and a correlation of 0.817 with frontal-to-posterior (F-P) traveling waves. Similarly, the relative phase averaged over parietal regions shows a correlation of 0.892 with directional wPLI over parietal regions, and a negative correlation of −0.813 with front-to-back traveling waves (all *p*-values *<* 0.001). Thus, the relative phase integrates two different approaches to reveal the directionality of inter-regional directionality. We also observed that the relative phase, traveling wave, and phase-based measures exhibit either no statistically significant correlations or only low correlations (*ρ ≤* 0.304) with amplitudes of the brain wave (see Supplementary Information Section B.1 for more detail), indicating that amplitude and phase dynamics contain distinct information [30].

### 2.2 Relative phase analysis across levels of consciousness

Monitoring the level of consciousness in general anesthesia is essential for ensuring patient safety and proper management. In this section, we investigate the relative phase dynamics in using the University of Michigan general anesthesia dataset [24, 29] to explore their relationship with consciousness levels. As each participant underwent general anesthesia through the administration of anesthetic agents (propofol followed by isoflurane), they gradually lost consciousness, progressing through distinct states: **eyes-closed resting state, loss of consciousness (LOC), burst period, suppression period**, and **recovery of consciousness (ROC)**. The transitions between loss and recovery of consciousness were determined by the participant’s ability to follow a verbal command to squeeze their hand. In essence, participants started in the resting state, then lost consciousness (LOC), entered deep anesthesia (burst and suppression periods), and finally regained consciousness (ROC). We note that the burst and suppression periods occur during the deepest stage of anesthesia, characterized by alternating bursts of brain activity (burst period) and suppressed activity (suppression period). (See Supplementary Information Section A.1 for the detailed anesthesia protocol.)

For each anesthesia state, we computed relative phase topographic map at each time point (Figures 1B and 1C; see Methods). Notably, the dynamics of relative phase patterns reveal distinct differences between conscious and unconscious states (Figure 2A). In the *eyes-closed* resting state, the front-to-back and back-to-front relative phase patterns repeat consistently over time. In contrast, the patterns become random and disorganized in the suppression state, the unconsciousness state occurring during the deepest anesthesia.

**Fig. 2.**
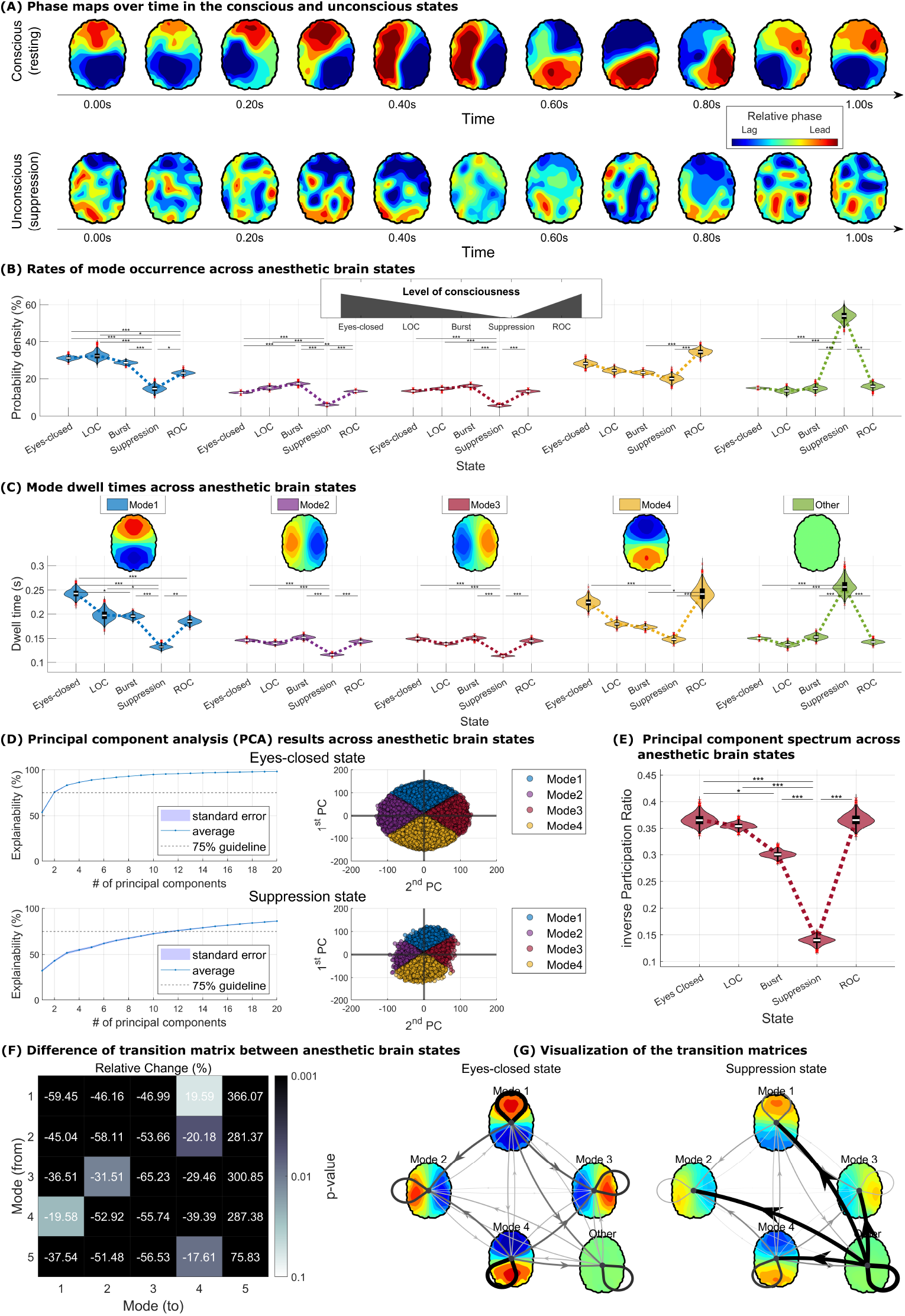
Statistics of relative phase dynamics in the general anesthesia dataset. (A) Representative topographic maps of relative phase snapshots during the *eyes-closed* resting state and the *suppression* state. The pattern becomes more random during the *suppression* state. To account for limited sample sizes, we performed 1,000 bootstrap resampling iterations in panels (B), (C), and (E). (B) Probability density functions of relative phase modes vary with the level of consciousness. Notably, modes 1 and 4 are strongly associated with consciousness depth, suggesting their potential as biomarkers for monitoring brain state. (C) Dwell time–defined as the duration sustained in each mode–also differs systematically across brain states. (D) Explainability in principal component analysis (PCA), reflecting how variance is distributed across components, shows distinct patterns: in the *eyes-closed* state, the first three principal components account for over 70% of total variance, whereas in the *suppression* state, at least 20 components are needed to reach a similar threshold. (E) The inverse Participation Ratio (iPR) decreases with deeper levels of unconsciousness, indicating reduced heterogeneity in explainability across components. To highlight the difference between the *eyes-closed* and *suppression* states, we present (F) the relative change (%) of the transition matrix between two states–representing how the suppression state differs from the *eyes-closed* state– and (G) the transition matrices visualized as transition networks, respectively. The relative changes were computed by subtracting the mean transition rates of the *eyes-closed* state from those of the *suppression* state, normalized by the *eyes-closed* state’s mean, and expressed as percentage change.

We further examined the relative phase properties, such as mode distribution and mode dwell time, to find the relationship between the relative phase and the level of consciousness. We applied *K*-means to identify dominant phase dynamics (Figure 1C; see Methods). This analysis revealed four distinct relative phase topographic patterns: two dominant modes, characterized by front-to-back and back-to-front patterns (frontal area phase-leading parietal area and vice-versa), and two transient modes, defined by left-to-right and right-to-left patterns (left hemisphere phase-leading right hemisphere and vice-versa) (Figures 2B and 2C).

Occupancy rates and dwell times revealed that, in the resting state, front-to-back (mode 1) and back-to-front (mode 4) patterns were predominant, with higher occupancy rates and prolonged dwell times of approximately 200 milliseconds. In contrast, left-to-right (mode 2) and right-to-left (mode 3) patterns were more transient, exhibiting lower occupancy rates and shorter dwell times (Figures 2B and 2C). Transition matrix analysis further demonstrated that mode transitions predominantly occurred through the left-to-right or right-to-left patterns, serving as transition modes between dominant modes of front-to-back and back-to-front patterns (Figure 5C, and Supplementary Section B.5).

For every time series of relative phase topographic maps, we can also apply linear regression taking the centroids of the *K*-means clustering as the vector of regressors, where the relative phase topographic map of each time point becomes the dependent variable ŷ(*t*) (Figure 1C). we applied linear regression taking the four centroids of the *K*-means clustering as the two vectors of regressors: Here, the first vector is a centroid representing front-to-back pattern (negative of which represent back-to-front pattern), and the second vector is a centroid representing left-to-right (negative of which represent right-to-left pattern) (see the regression formula in Figure 1C). By observing how the coefficients for the first and second vector (*β*_1_(*t*) and *β*_2_(*t*) respectively) change over time, we can quantify the dominance of each modes, and also cluster topographic map from each time point *t* into the one of four modes. In other words, linear regression is used to quantify the contribution of these two paired modes at every time point: if |*β*_1_(*t*)| *>* |*β*_2_(*t*)|, the topographic map at time *t* is either in front-to-back mode (if *β*_1_(*t*) *>* 0) or back-to-front mode (*β*_1_(*t*) *<* 0); if |*β*_1_(*t*)| *<* |*β*_2_(*t*)|, the topographic map at time *t*is either in left-to-right mode (if *β*_2_(*t*) *>* 0) or right-to-left mode (*β*_2_(*t*) *<* 0). We note that such linear regression approach is mathematically equivalent to principal component analysis (PCA), and the resulting clustering closely aligns with the *K*-means clustering results (see Supplementary Figure S6). This equivalence arises from the inherent relationship between *K*-means clustering and PCA, where each principal component closely corresponds to two paired modes identified by *K*-means clustering. Front-to-back and back-to-front modes varied systematically across levels of consciousness (Figures 2B and 2C). Additionally, we introduced the category ‘Other’ for topographic maps which are difficult to classified to either of the four modes: A topographic map at time point *t* is labeled as ‘Other’ when the regression residual *ε*(*t*) at *t* exceeds the 85th percentile (top 15%) of all residuals across time points, indicating that the topographic map at *t* is not well characterized by *β*_1_(*t*) and *β*_2_(*t*). (Figures S8 and S9 in the Supplementary Information).

In summary, the occupancy rate and dwell time of front-to-back mode (mode 1) and back-to-front mode (mode 4) tended to decrease as consciousness diminished, then increased to levels comparable to the resting state upon recovery. The occupancy rate of mode 1 changed from 31.02% (resting) to 32.40% (LOC), 28.48% (Burst), 14.64% (Suppression), and 23.01% (ROC). Mode 4 shifted from 28.02% (resting) to 24.31% (LOC), 23.22% (Burst), 20.03% (Suppression), and 34.49% (ROC). Statistically significant differences in mode 1 occupancy rates were observed between resting and suppression states, resting and ROC states, LOC and suppression states, LOC and ROC states, burst and suppression states, and suppression and ROC states (one-way ANOVA test performed for all comparisons; see Methods; all *p*-values *<* 0.05). For mode 4, significant differences occurred between burst and ROC states, and suppression and ROC states (all *p*-values*<* 0.05).

Mode 1 dwell time changed from 242.7 ms (resting) to 197.4 ms (LOC), 195.5 ms (Burst), 132.2 ms (Suppression), and 185.5 ms (ROC). Mode 4 dwell time varied from 224.7 ms (resting) to 179.4 ms (LOC), 173.1 ms (Burst), 148.0 ms (Suppression), and 242.8 ms (ROC). Significant differences in mode 1 dwell time were found between resting and burst states, resting and suppression states, resting and ROC states, LOC and suppression states, burst and suppression states, and suppression and ROC states (all *p*-values *<* 0.05). For mode 4, significant differences were observed between resting and suppression states, LOC and suppression states, burst and suppression states, and suppression and ROC states (all *p*-values *<* 0.001).

Our findings from the regression model align with those obtained through principal component analysis (PCA). As shown in Figures 2D and 2E, the inverse participation ratio (iPR)–which reflects how many principal components are required to explain the given data–decreased as consciousness diminished. Specifically, iPR changed from 36.59% (resting) to 35.45% (LOC), 30.07% (Burst), and 13.95% (Suppression), before returning to 36.61% (ROC). Significant differences in iPR were observed between resting and burst states, resting and suppression states, LOC and suppression states, burst and suppression states, and suppression and ROC (all *p*-values *<* 0.05). These results confirm, using PCA, that relative phase patterns became more random as consciousness decreased.

Furthermore, we examined the transition properties among the modes. In addition to mode distribution and dwell time, transition patterns also differed between conscious and unconscious states (Figures 2F and 2G). Specifically, transition rates into the ‘Other’ mode were significantly higher in the suppression state compared to the eyes-closed state, indicating greater randomness in brain wave patterns. In contrast, transitions among modes 1, 2, 3, and 4 were generally reduced in the suppression state.

In summary, our findings suggest that properties of the relative phase patterns, particularly the ‘front-to-back’ and ‘back-to-front’ modes, serve as reliable markers for distinguishing different states of consciousness.

### 2.3 Relative phase analysis of simultaneous EEG–fMRI during resting and stimulus states

To further examine the functional significance of front-to-back and back-to-front phase modes, we compared their expression with the activity levels in well-established functional brain networks derived from fMRI BOLD signals. This analysis was performed using a simultaneous EEG–fMRI dataset, which included alternating periods of eyes-closed resting state and a flashing checkerboard visual task [25]. The task consisted of 20-second blocks of eyes-closed rest and 12 Hz flashing checkerboard stimulation, a paradigm known to robustly activate the visual network (VN).

We confirmed that the visual network exhibited increased activation during the checkerboard stimulation periods and decreased activation during rest. Differences in BOLD activity between the rest and stimulation conditions are shown in Figure 3A, revealing that higher-order cognitive regions were more active during rest, whereas visual sensory areas showed greater activation during stimulation. For this analysis, BOLD signals were first averaged across voxels within each region defined by the 1,000-parcel Schaefer atlas [31], and subsequently Z-scored to normalize voxel-specific signal variability (see Methods for details). The Z-scored time series were then averaged across trials and participants to derive group-level time series. Finally, the difference between group-averaged signals during rest and stimulation periods was computed.

**Fig. 3.**
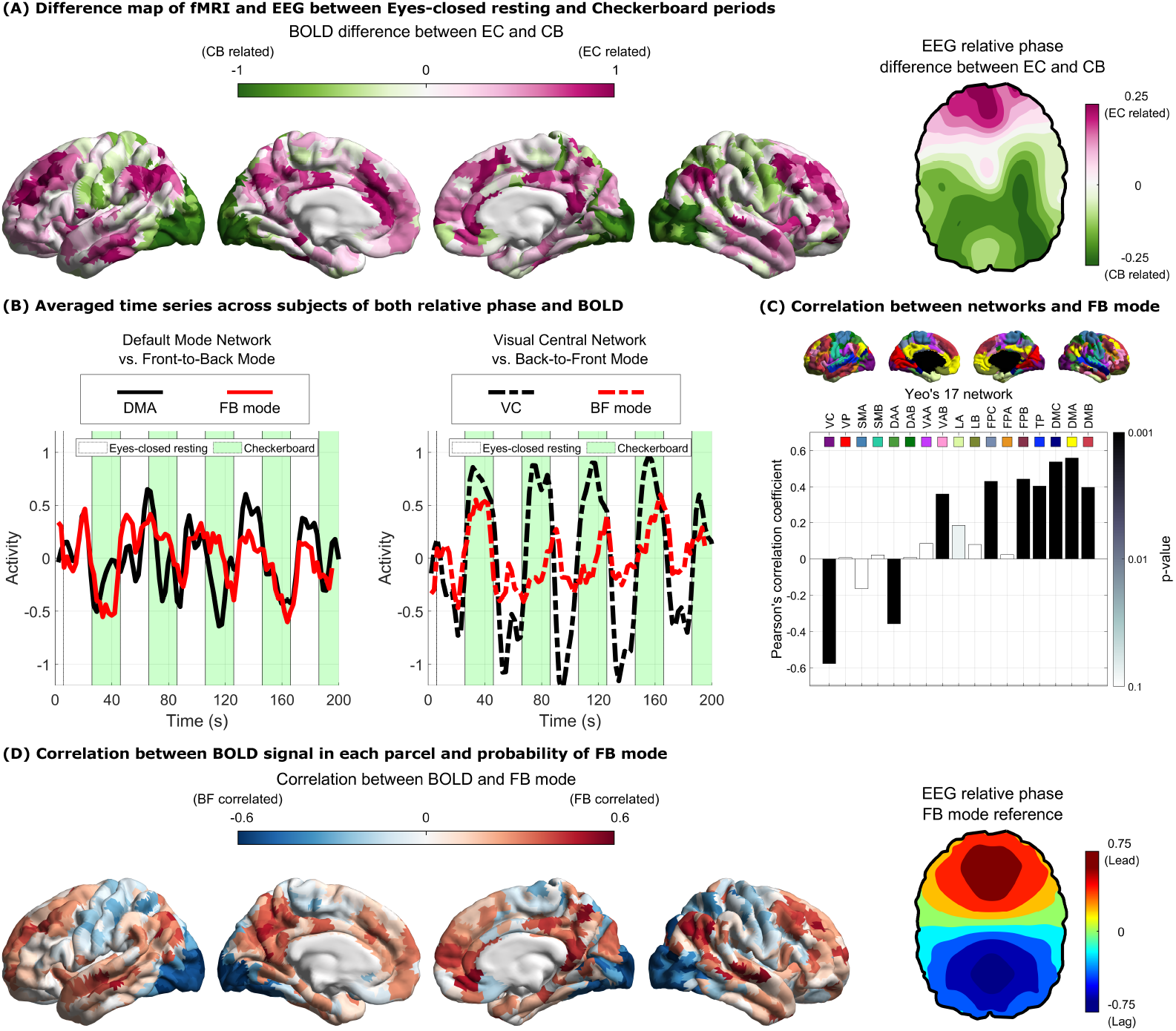
Brain network correspondence of relative phase patterns using simultaneous EEG–fMRI recordings. To investigate the functional relevance of front-to-back (FB) and back-to-front (BF) modes identified through EEG-based Relative Phase Analysis (RPA), we compared these patterns with BOLD activity measured via fMRI during two conditions: eyes-closed resting state (EC) and a 12 Hz flashing checkerboard task (CB). (A) Differences between EC and CB states are shown for both cortical BOLD surface maps and EEG relative phase topographic maps. In the BOLD maps, purple regions indicate greater activity during EC, and green regions indicate greater activity during CB. In the EEG relative phase topographic maps, purple regions represent phase-leading areas during EC, while green regions indicate phase-leading areas during CB. (B) Time series of EEG-derived *β*_1_ values (where positive values reflect FB mode) and BOLD activity from the Default Mode A (DMA) network are shown on the left; the inverse, *−β*_1_ (representing BF mode), is plotted alongside activity from the Visual Central (VC) network on the right. *β*_1_ is positively correlated with DMA activity and negatively correlated with VC activity. (C) Pearson’s correlation coefficients between *β*_1_ and all Yeo’s 17 networks. (D) Visualization of Pearson’s correlation coefficients between *β*_1_ and BOLD activities (Scheafer’s 1,000 parcellation). The red colored areas are highly correlated with FB mode and blue colored areas are highly correlated with BF mode. The acronyms for Yeo’s 17 networks: VC-Visual Central, CP-Visual Peripheral, SMA-Somatomotor A, SMB-Somatomotor B, DAA-Dorsal Attention A, DAB-Dorsal Attention B, VAA-Ventral Attention A, VAB-Ventral Attention B, LA-Limbic A, LB-Limbic B, FPC-Frontoparietal C, FPA-Frontoparietal A, FPB-Frontoparietal B, TP-Temporal Parietal, DMC-Default C, DMA-Default A, DMB-Default B.

To compare EEG phase dynamics across conditions, relative phase topographic maps were first averaged across trials and then across participants. The difference between the resulting group-level maps for the rest and stimulation periods was then computed. We observed that the phase-leading pattern in frontal regions was more prominent during the resting periods, whereas the phase-leading pattern in parietal regions was more pronounced during the checkerboard stimulation (Figure 3A, right). From a time-series perspective, the long-term trend—persisting after averaging across trials and participants—revealed a systematic bias toward either the front-to-back or back-to-front mode, depending on the brain state.

To more comprehensively characterize the relationship between EEG relative phase patterns dynamics and fMRI BOLD signals, we examined activation across 17 canonical functional networks using established parcellations [32]. We observed a strong positive correlation between the occurrence of the front-to-back (FB) mode—quantified by the regression coefficient *β*_1_(*t*)–and activity of the Default Mode Network (DMN): DMA, DMB, and DMC (all *r >* 0.5, *p <* 0.001). Conversely, the back-to-front (BF) mode—represented by negative of *β*_1_(*t*)–was positively correlated with activity in the Visual Central (VC) network (*r* = 0.58, *p <* 0.001) (Figures 3B and 3C). To account for delayed hemodynamic and electrophysiological responses, event onsets were shifted by 6 seconds–effectively aligning event boundaries 6 seconds later relative to the recorded BOLD and EEG signals. Notably, the cross-correlation between BOLD activity and EEG relative phase dynamics (represented by *β*_1_) peaked at zero lag, indicating a strong temporal correspondence between the two signals. This suggests that EEG relative phase dynamics are not only correlated with BOLD signals but may also exhibit similar delays relative to event onsets, reflecting shared latency in their response to external stimuli. We note that, for consistency, the same 6-second shift in event onsets was also applied to the earlier EEG and BOLD comparisons between resting and stimulation periods (Figure 3A).

These results suggest that posterior-to-anterior directionality (BF mode) is associated with externally oriented sensory processing such as VN, whereas anterior-to-posterior directionality (FB mode) align with internally directed, higher-order processing such as the DMN. To further validate this association, we performed whole-brain temporal correlation mapping between front-to-back mode expression—again represented by *β*_1_–and BOLD signals across the 1,000-parcel Schaefer atlas. Specifically, we computed temporal correlations between *β*_1_ and the mean BOLD signal within each of the 1,000 parcels (Figure 3D). This analysis revealed that front-to-back mode expression was not only strongly correlated with the DMN but also with other higher-order association networks, including the Frontoparietal Network (FPN; FPC and FPB) and the Temporal-Parietal (TP) network, all significant at *p <* 0.001 (Figure 3C). Together, these findings indicate that anterior-to-posterior directionality reflects engagement of higher-order cortical networks involved in internally guided cognitive processes.

Based on these results, we define the front-to-back pattern as the **top-down** mode, representing information flow from higher-order cognitive anterior regions to posterior regions. Conversely, we designate the back-to-front pattern as the **bottom-up** mode, representing information flows in the opposite direction.

### 2.4 Relative phase analysis in ADHD vs. Control groups

Using simultaneous EEG–fMRI data, we demonstrated that the fluctuations captured by Relative Phase Analysis (RPA) likely reflect the activity and interactions of functional brain networks. To further validate the functional significance of dominant phase modes and to explore the cognitive relevance of this approach, we applied RPA to resting-state EEG data from individuals with the ADHD-inattentive subtype and typically developing controls.

There are multiple neuroscientific approaches to understanding ADHD symptoms, ranging from neurotransmitter-level theories to network- and cognitive-level models [13, 33–35]. Among these, cognitive-level theories suggest that individuals with ADHD exhibit reduced top-down signaling compared to controls, along with impaired coordination between top-down and bottom-up processes [26, 36, 37]. This imbalance is believed to contribute to the inattentive symptoms commonly observed in ADHD. In the previous section, we showed that RPA captures fluctuations between top-down (front-to-back) and bottom-up (back-to-front) modes by correlating EEG relative phase dynamics with simultaneously acquired fMRI signals on a matched time scale. Building on this, we applied the relative phase measure to EEG data from both ADHD-inattentive and control groups to assess whether our findings align with previous research on attentional impairments in ADHD. This approach may provide cognitive-level evidence that relative phase patterns reflect attentional processing as a key component of conscious function.

Moreover, we aim to explore the potential of relative phase dynamics as a biomarker for diagnosing the ADHD-inattentive subtype. Currently, the theta/beta ratio (TBR) is the only EEG-based biomarker approved by the U.S. Food and Drug Administration (FDA) for ADHD diagnosis. However, TBR has shown considerable variability across studies and is most effective in identifying the ADHD combined subtype, which involves both inattentive and hyperactive-impulsive symptoms [38, 39]. By comparing RPA with conventional measures, we seek to assess whether relative phase analysis can serve as a more reliable diagnostic tool for ADHD-inattentive presentations.

We analyzed EEG resting-state data from the ADHD-inattentive and control groups in the Healthy Brain Network (HBN) data released by the Child Mind Institute [40]. Following the same methodology as before, we conducted 4-means clustering and calculated both the mode occupancy rate and the dwell time in each of the four modes. The topological dynamics in the eyes-closed resting state derived from relative phase over time illustrate the differences between the ADHD and Control groups (Figure 4A). The Control group shows continuous changes at ∼0.2-second intervals, whereas the ADHD group tends to stay in the back-to-front (bottom-up) mode for longer periods. These topological dynamics alone suggest that the RPA displays an imbalanced pattern in the ADHD group.

**Fig. 4.**
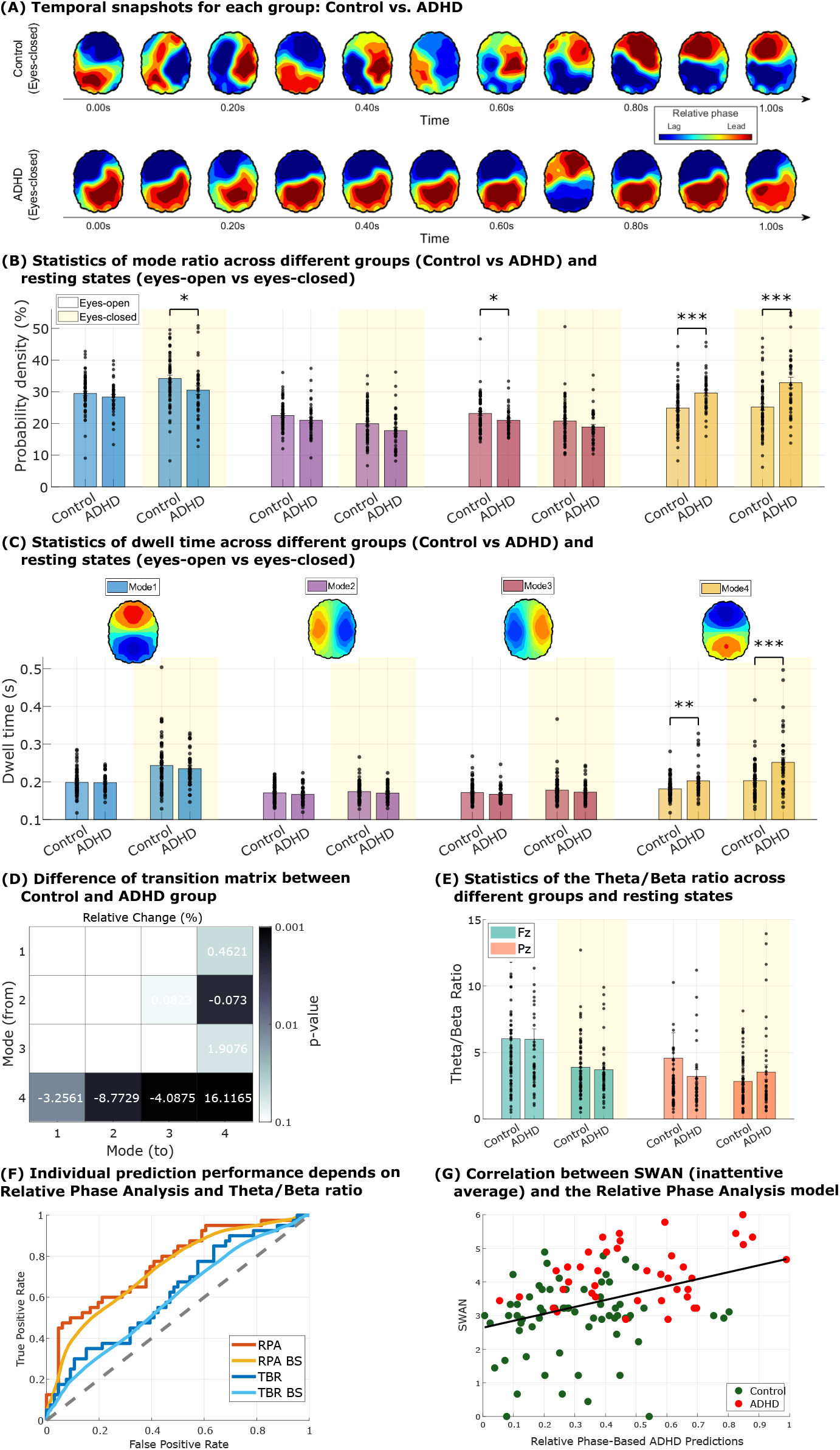
Relative Phase Analysis applied to the HBN ADHD-inattentive dataset. (A) Representative snapshots of relative phase topographic maps during the eyes-closed resting state for the control group and the ADHD-inattentive group. ADHD-inattentive group dwell on back-to-front mode more. (B) Probability density functions for both groups, shown across eyes-open (white background) and eyes-closed (yellow background) resting states. Asterisks indicate significance levels from two-sample t-tests: \**p* ≤ 0.05, * * *p* ≤ 0.01, and* * * *p* ≤ 0.001. (C) Dwell time distributions across relative phase modes for the control and ADHD-inattentive groups. (D) Relative difference matrix between the transition matrices of the two groups during the eyes-closed state. Differences were computed by subtracting the mean transition rates of the control group from those of the ADHD group, normalized by the control group’s mean, and expressed as percentage change. (E) Theta/beta ratios at the Fz and Pz electrodes for both groups. Green and orange bars represent the Fz and Pz channels, respectively. (F) Receiver Operating Characteristic (ROC) curves for individual-level diagnostic prediction using Relative Phase Analysis and the theta/beta ratio. (G) Correlation between SWAN inattentive scores and predicted diagnostic values from the Relative Phase logistic regression model.

**Fig. 5.**
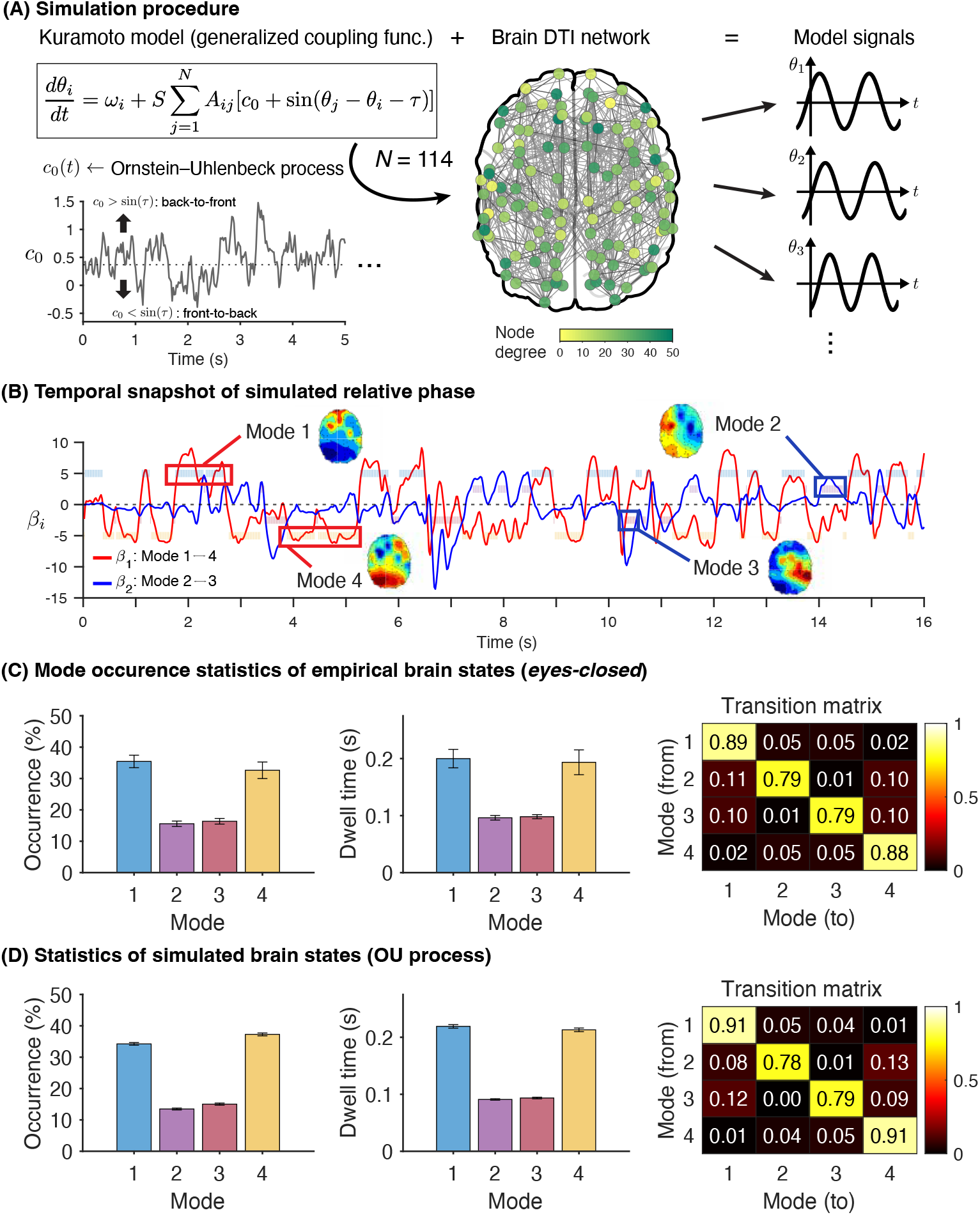
Simulation of relative phase dynamics using the Kuramoto coupled oscillator model. (A) Schematic of the simulation process using the Kuramoto model with a generalized coupling function on the human brain network. The *c*_0_ parameter in the model is time-dependent and evolves according to the Ornstein-Uhlenbeck stochastic process, as shown in the example trajectory of *c*_0_. Horizontal dotted lines indicate the fixed point at *c*_0_ = sin (*τ*). The model generates phase signals for each brain region (*N* = 114). (B) Snapshot of relative phase topographic map from model-generated signals. (C) Statistics of empirical mode distributions in the University of Michigan (UofM) general anesthesia dataset in the conscious state (*eyes-closed* resting state). Plotted are the distribution of occurrences (left), dwell time (middle), and transition matrices among the four brain states (right). Error bars = s.e.m. across subjects. (D) Same plot as in C but for simulated model data, averaged from 50 unique initial conditions. Simulation parameters: *S* = 1.8, *τ* = 0.12*π, ψ* = 6, *σ* = 1.5. Error bars = s.e.m. across simulation instances.

Analysis of the mode distribution and mode dwell time quantitatively demonstrated these group differences (Figures 4B and 4C). In both eyes-closed and eyes-open resting states, the ADHD-inattentive subtype group exhibited significantly greater rates of mode occupancy and greater dwell times in mode 4 (back-to-front or bottom-up mode) compared to the control group. In the eyes-closed state, the differences between the two groups were 30.6% in mode distribution and 23.9% in dwell time, relative to the control group, and these differences were significant in both measures (two-sample t-test, *p <* 0.001), and these results remained significant after applying the more stringent False Discovery Rate (FDR) correction. Thus, the differences in the posterior-to-anterior directionality mode could reliably distinguish the ADHD-inattentive group from the Control group. Furthermore, an analysis of probability density of mode 1 (front-to-back or top-down mode) in the eyes-closed condition reveals that the ADHD group shows a lower value than the control group at a 95% confidence level (*p* = 0.036, Figure 4C). This finding aligns with bias towards mode 4, indicating a deficiency in top-down signaling of ADHD-inattentive group.

We also analyzed the between-mode transition matrix for each group, and computed the difference in between-mode transition probabilities for the ADHD group and Control group values (Figure 4D). We found that participants in the ADHD group exhibited greater mode 4 to mode 4 transitions (i.e. staying in mode 4; *p <* 0.001). This imbalance between mode 1 (front-to-back, top-down) and mode 4 (back-to-front, bottom-up), along with slower fluctuations, supports the current hypothesis regarding ADHD inattention symptoms.

To compare measures, we calculated the TBR using the same HBN dataset (Figure 4E). The TBR is usually calculated by obtaining power values from the Cz, Fz, and Pz channels to assess differences between the two groups. However, since the HBN dataset uses Cz as the reference channel, our study calculated solely on the TBR values from the Fz and Pz channels. Previous research suggests that the differences between these two channels and the Cz channel are minimal [41]. Our analysis of the HBN data indicated no significant TBR differences between the control group and the ADHD-inattentive group, reaffirming findings from earlier studies [38, 39]. These results suggest that the relative phase pattern is a more effective biomarker for the ADHD-inattentive subtype.

Receiver Operating Characteristic (ROC) curve analysis showed that the model based on Relative Phase Analysis (RPA) outperformed the theta/beta ratio (TBR) model in individual-level diagnostic prediction (Figure 4F). In our model, we used logistic regression to determine the weights, with the diagnostic outcome as the dependent variable and the independent variables being the occurrence of mode 4, the dwell time in mode 4, and the self-transition probability from mode 4 to mode 4. We then used these weights to calculate an assessment score for each subject based on their independent variables. The RPA model achieved a performance score of 0.76, and 0.73 when validated via bootstrap method, whereas the TBR model yielded scores of 0.62 and 0.59, respectively.

Lastly, we examined the correlation between the RPA model’s diagnostic prediction score and the average of Strengths and Weaknesses of ADHD Symptoms and Normal Behavior Scale (SWAN) inattentive score [42] with the correlation coefficient as 0.38 (*p <* 0.001). This result indicates a meaningful association between the RPA model and behavioral measures (Figure 4G). While the SWAN score reduces the likelihood of false negatives in identifying ADHD, the RPA model helps reduce false positives. Notably, distinguishing between control and ADHD subjects in the upper-left quadrant may offer valuable insights, warranting further investigation from this perspective in future studies.

These analyses confirm that the relative phase dynamics are consistent with the existing hypothesis about ADHD-inattentive symptoms: a deficiency in top-down mode and an imbalance in communication between top-down and bottom-up modes. This finding strengthens our conclusion that the relative phase pattern can be linked to top-down and bottom-up directionalities in the brain, and highlights the importance of the relative phase pattern and its connection to ADHD-inattentive symptoms. Additionally, the comparison with the TBR measure indicates that RPA could serve as a biomarker for ADHD-inattentive subtype.

### 2.5 Simulating relative phase dynamics via coupled-oscillator model

Finally, to elucidate the mechanism underlying the mode fluctuations, we ran simulations of relative phase dynamics using a Kuramoto model, a well-known canonical coupled oscillator model frequently used to model large-scale cortical dynamics. Specifically, we utilized a Kuramoto model with a generalized coupling function that includes a constant term (*c*_0_), which results in a more realistic first order approximation of many coupling functions obtained from the phase reduction method [27, 28, 43]. This term is also naturally derived from a control parameter that represents an intermediary between the diffusive and direct coupling schemes in limit-cycle oscillators [44, 45]. Prior theoretical work has shown that inclusion of this parameter in the coupling function results in a rich repertoire of phase dynamics, especially relevant to the transitions between two distinct and dominant modes in the brain: one in which high-degree cortical regions phase-lag the network becoming sink of the directionality, and one in which the hub regions phase-lead the network and becoming source of the directionality [45]. Motivated by this finding, we hypothesized that the relative phase dynamics can be simulated with random fluctuations in this *c*_0_ parameter, which is linked to the phase-lead/lag relationship of the higher-degree regions and the resulting front-to-back/back-to-front directionality (see Supplementary Figure S25 for more details). We simulated the Kuramoto model on a human structural cortical network with *N* = 114 regions [46], with each node generating a phase signal according to the dynamics of the model (Figure 5A). The fluctuations in *c*_0_ were modelled with the Ornstein-Uhlenbeck (OU) process. Using grid search, we then estimated the parameters of the stochastic process that best match the empirical mode statistics, focusing on the UofM general anesthesia data (conscious, eyes-closed).

By applying the same procedure for the empirical data, we obtained the regression weights for each time point (*β*_1_ and *β*_2_) specifying the mode distribution over different brain states. Example temporal snapshots of simulated topographic plot for each mode is illustrated in Figure 5B. In particular, we found that mode 1 was indeed associated with high-degree nodes phase-lagging relative to the system, while mode 4 showed the opposite effect where high-degree nodes were phase-leading (Supplementary Figure S25D). This was reflected as significant correlations between the averaged relative phase and the anterior-posterior coordinates of the brain regions (Supplementary Figure S25E), consistent with the regression weights specifying back-to-front directionality (*β*_1_). More specifically, when the brain state occupied mode 1, the correlation was negative (*r* = −0.249, *p* = 0.00752) such that the more anterior regions tended to phase-lead the system (Supplementary Figure S25E, middle). In contrast, the correlation was positive (*r* = 0.255, *p* = 0.00608) during mode 4, in which the more posterior regions tended to phase-lead the system (Supplementary Figure S25E, right).

By inspecting the statistics of the simulated mode occurrences and dwell time, we further found that the Kuramoto model captured the key characteristics of the empirical data in Figure 5C. As explained earlier, the empirical mode fluctuations during conscious resting state exhibited a pattern where modes 1 and 4 occurred more dominantly than modes 2 and 3 (Figure 5C, left). This pattern is also reflected in the dwell time distribution, in which modes 1 and 4 are occupied for a longer period of time than modes 2 and 3 on average (Figure 5C, middle). From the transition probability matrix (Figure 5C, right), the brain states tended to stay in the same mode (high transition rates along the diagonal), while modes 2 and 3 served as the transient states between modes 1 and 4. (For example, mode 2 or 3 was more likely to transition into mode 1 or 4 than transitioning into 3 or 2.) These characteristics were well reflected in the simulated relative phase dynamics in Figure 5D. That is, the simulated modes fluctuations showed the dominance of modes 1 and 4 compared to modes 2 and 3 in terms of both occurrence rate and dwell time (Figure 5C, left and middle). The simulated transition probability matrix (Figure 5D, right) also highlighted the key tendency reflected in the empirical data, wherein the transitions between the modes 2 and 3 were near zero and instead they were transiently visited from 1 and 4.

Overall, these results illustrate that a canonical model of coupled oscillators, such as the Kuramoto model equipped with a simple stochastic process in just one parameter, can replicate the key characteristics of the relative phase dynamics in the human brain. The model simulations may also point to the universality of the relative phase dynamics observed in the human brain network. We note that the OU process is not the only stochastic process that can replicate the empirical data; we found that other processes, involving more than one stable fixed point, were also able to capture the data for a specific range of parameters (Supplementary Figures S26D and S26E). These findings raise the possibility that the observed top-down and bottom-up modes could be understood as the attractor states in a network of coupled brain regions.

## 3 Discussion

What are the dominant modes of directionality at the whole-brain level? What are the real-time transition dynamics between these modes? We addressed these questions by establishing relative phase as a robust, versatile, and clinically relevant measure of brain dynamics. Leveraging RPA, we found that: (i) the anterior-to-posterior (top-down) versus posterior-to-anterior (bottom-up) directional patterns are fundamental modes of human brain dynamics, with the brain rapidly fluctuating between these two states on a sub-second scale; (ii) these dynamic properties are most pronounced during full consciousness and diminish as consciousness fades; (iii) the directional modes are associated with the BOLD activity of corresponding functional networks (anterior-to-posterior mode with DMN and FPN, posterior-to-anterior mode with VN); (iv) developing individuals and those with ADHD exhibit group-level differences in their relative-phase profiles and dynamics; and (v) a generalized coupled-oscillator brain network model reproduces the observed mode transition dynamics.

### Relative Phase Analysis Reveals a Real-Time, Whole-Brain Dynamics of Sub-Second Anterior-Posterior Directionality Fluctuations

Relative phase analysis (RPA) provides a real-time measure in quantifying phase dynamics, enhancing our ability to investigate large-scale neural interactions beyond previous methods. Unlike time-window-based phase-lag measures or region-specific traveling-wave analyses, RPA offers continuous tracking of neural phase relationships across the whole cerebral cortex. As demonstrated in the Results section, the method can still incorporate time-window averaging and region-specific calculations to examine long-term and spatially localized dynamics. However, its real-time nature offers a significant advantage for investigating fine temporal dynamics. Additionally, RPA holds promise for neurofeedback applications, where the ability to monitor real-time neural dynamics is crucial.

Leveraging RPA, we identified sub-second transitions between dominant top-down and bottom-up directional modes. Prior studies suggest that such fluctuations, or more generally, the alternation between internally and externally biased processing states, occur across multiple spatial and temporal scales [17]. This oscillatory balance has even been proposed as a fundamental signature of human brain information processing across multiple levels of organization [17]. Event-related-potential (ERP) studies from EEG and ECoG indicate that the brain’s response to external stimuli occurs within sub-second timescales, suggesting that bottom-up vs. top-down switching may operate within a similar temporal window. Our findings confirm this, demonstrating that these switching occurs approximately every 200 ms, spanning the whole brain in a dynamic interplay.

Crucially, we observed that these fluctuations are not solely stimulus-driven but also emerge spontaneously during resting states, suggesting an intrinsic top-down vs. bottom-up modulation independent of external input. This finding highlights a fundamental, self-organizing mechanism of cortical dynamics. For our future studies, we aim to leverage RPA in ERP paradigms, exploring its potential for ERD analysis and its role in cognitive function. Additionally, we observed that both top-down and bottom-up directionalities extend across all frequency bands up to 50 Hz. Future investigations will examine whether the balance between top-down and bottom-up processing varies across frequency bands in response to different stimuli, potentially uncovering frequency-dependent mechanisms of neural information flow [9, 47, 48].

### Sub-Second Anterior-Posterior Directionality Fluctuations Link to BOLD Activity Dynamics and Distinguish Diverse States of Consciousness and ADHD

Our relative phase analysis may offer a new approach for assessing consciousness levels in clinical settings. Distinguishing brain states across different levels of consciousness remains a fundamental challenge, particularly in clinical applications such as general anesthesia. While phase- and amplitude-based measures have shown partial success in identifying consciousness levels, they do not fully capture all aspects of anesthesia. For instance, the amplitude dynamics of burst states during anesthesia closely resemble those of resting states, making differentiation difficult for some conventional measures. In contrast, our analysis of the distribution of top-down and bottom-up modes effectively characterizes variations in consciousness levels and successfully detects the reduction in burst states. This finding further supports the measure’s ability to quantify reductions in dominant directional modes, which we propose as a key mechanism underlying full consciousness. Notably, the decrease in dominant modes correlated with consciousness levels occurred also in the low beta band as well as the alpha band. Further exploration of how different anesthetic agents modulate mode distributions across frequency bands could provide deeper insights into comparative anesthetic mechanisms and their effects on neural dynamics [1, 4].

Modes 1 and 4, associated respectively with top-down and bottom-up neural processes, typically exhibit dwell times of around 200 ms during conscious states. This duration closely aligns with the 200–300 msec window identified by Global Neuronal Workspace Theory (GNWT) as critical for conscious awareness [49–51]. Under anesthesia, however, dwell times are substantially reduced: mode 1 decreases from approximately 243 ms (during eyes-closed resting) to 132 ms (during unconscious suppression), while mode 4 shortens from about 225 ms to 148 ms. GNWT proposes that conscious perception (global ignition) requires approximately 200 ms of sustained, coherent neural activity. Thus, our findings suggest that dwell times below this critical time duration impair the brain’s ability to achieve global ignition, thereby disrupting conscious awareness. These results align with GNWT predictions, reinforcing that the approximately ∼200 ms dwell time observed in our study represents a neural dynamic essential for consciousness. Notably, in contrast to previous studies–which suggest anesthesia induces unconsciousness mainly by selectively suppressing long-latency signals associated with higher-order cognitive processes– our results indicate a different mechanism [52]. Our results suggest that anesthesia significantly alters brain dynamics by shortening dwell times of both top-down and bottom-up modes, resulting in insufficient neural coherence duration–shorter than the critical time duration of ~200 ms– to effectively integrate sensory inputs into conscious perception.

Simultaneous EEG–fMRI not only bridges brain wave activity with BOLD signals but also uncovers insights unattainable through a single modality. Here, we establish a link between directional mode transitions in brain waves and corresponding functional network transitions in BOLD activity. Using a flashing checkerboard stimulus, a well-established paradigm for eliciting reliable BOLD activation patterns, we interpret anterior-to-posterior and posterior-to-anterior directionality as reflecting top-down and bottom-up processing, respectively. Future research will extend this analysis to EEG–fMRI data acquired with more nuanced and naturalistic stimuli, aiming to capture mode transition dynamics in more natural settings. Additionally, applying relative phase analysis (RPA) directly to BOLD time series may provide novel insights into the directional propagation of BOLD signals, complementing recent methodological advances. A systematic comparison and integration of RPA with established measures could ultimately lead to a more robust framework for inferring BOLD dynamics, overcoming the inherent temporal resolution limitations of fMRI.

Our RPA approach successfully distinguishes between the inattentive type ADHD group and the typically developed control group demonstrating high performance in individual diagnostic prediction. These findings suggest the potential validity of RPA as a new EEG biomarker for ADHD, supplementing the TBR measure which remains a topic of debate regarding its effectiveness in diagnosing ADHD. Furthermore, the observed increase in bottom-up mode and decrease in top-down mode aligns with previous theories suggesting that attention deficit symptoms are associated with weakened top-down inhibition of bottom-up processes [26]. Considering the existing gap between cognitive-level explanations of ADHD and brain imaging measures, our results support the role of RPA in elucidating the neuroscientific mechanisms underlying ADHD-inattentive symptoms. By identifying group-level differences in resting-state brain dynamics, our study connects with previous research showing that top-down inhibition is less active in ADHD individuals in the absence of cognitively demanding tasks [26, 53]. This suggests that RPA could serve as a neuroscientific approach within integrative models of ADHD, which incorporate cognition, emotion, and environmental factors [35, 54, 55]. Furthermore, the measure may offer the potential for developing more targeted biomarkers and interventions, as the validity of such interventions may be measured with the changes in the RPA. Therefore, we expect that applying RPA to ADHD and other neuro-disorders will provide valuable insights into their underlying mechanisms. In the future, developing more sophisticated AI models for individual predictions and investigating the relationship between the RPA model and other behavioral measures could enhance the potential of relative phase patterns for diagnosis and symptom reduction in clinical settings.

In this study, we primarily focused on validating the differences in top-down and bottom-up directionality across different groups, restricting the application of relative phase measures to the inattentive type of ADHD. This decision was influenced by the relevance of inattention symptoms to top-down mode activity and the sample size distribution within the HBN dataset. Additionally, to facilitate a clear comparison with the typically developed control group, we analyzed only adolescents aged 11 and older diagnosed solely with the inattentive subtype of ADHD. The age restriction is due to significant differences in EEG alpha spectrum peak that exist between individuals below and above the age of 10 [56]. In future studies, we aim to expand our analysis to hyperactive and combined types of ADHD, as well as comorbid conditions, to explore mode differences across these subgroups. The distinct findings among different groups will contribute to a deeper understanding of the underlying mechanisms of various neuro-symptoms.

### Computational Modeling of Sub-Second Directionality Modulation Provides Predictive Insights

Our computational modeling work suggests that the dominant modes and the transitions between them can occur spontaneously in a network of interacting neural masses. Namely, the anterior-posterior directionality and posterior-anterior directionality naturally occur due to their underlying brain network structure. With the addition of the stochastic coupling term (*c*_0_), the model exhibits the dominant modes as the attractors. In our modeling framework with coupled oscillators, the *c*_0_ parameter naturally arose as the first order approximation of many realistic coupling functions in the phase-reduced Kuramoto model [27, 28, 57]. In particular, this parameter serves as an intermediary term bridging the direct and diffusive coupling schemes [44, 45], which are two commonly assumed forms of coupling between oscillators. In the brain, these are often associated with the two major forms of coupling: electrical and chemical coupling of synapses, respectively [58, 59]. In our model, higher values of *c*_0_ (bottom-up mode) are related to the dominance of electric synapses in which neurons are coupled based on the difference in potentials through gap junctions, whereas lower values of *c*_0_ (top-down mode) are related to chemical synapses in which the generated action potentials are largely independent of potential difference between neurons. Thus our framework predicts that the transitions among the dominant modes reflect the changing ratios of these two forms of coupling in the large-scale brain network, which remain to be investigated empirically. Another possible interpretation of our model is that the *c*_0_ term reflects the relative dominance of external stimuli (sensory input) in the network, such that when c0 exceeds the sine function (sin (*τ*); Supplementary Figure S25A), external stimuli are more influential on the network dynamics, making the brain more receptive and responsive to sensory information (bottom-up processing mode). Conversely, when *c*_0_ is smaller than the sine function, intrinsic network dynamics are more influential, making the brain able to coordinate external sensory inputs with spontaneous internal brain rhythms—representing effective top-down modulation. Overall, our framework extends the previous theoretical proposals on the functional implications of having two co-existing forms of coupling in the oscillator system [44, 45, 58, 60, 61].

### Limitations and Future Perspectives

Finally, we mention some of the limitations of the current study, which we aim to address in future work i) Our analyses focused on resting-state data and a very strong external stimulus of flashing checkerboard. Extending RPA to task-based paradigms will be necessary to study the relationship between directionality modes and cognitive processes such as attention and memory. Establishing links between relative phase modes and specific behavioral or cognitive tasks will be in order for understanding the functional significance of top-down and bottom-up dynamics. For example, previous study [62] have reported a clear lag in the alpha oscillation in the posterior regions prior to strong distractions during a working memory task. Our RPA framework is consistent with this finding and could interpret the result as an increase in top-down control (i.e., mode 1) when distractors are expected. Future studies could probe whether such task-induced effects are more clearly demarcated by considering the global phase lead-/lag relationship in the whole brain. Additionally, auditory-dominant stimuli–such as those employed in the oddball paradigm [63–65]–may evoke distinct relative phase patterns than the back-to-front mode (mode 4), for auditory cortex is located at the temporal part of the cortex. This possibility will be investigated in future studies. ii) The relationship between relative phase modes and physiological signals or actions– such as electrocardiogram (ECG) activity, eye movements, or muscle activity–could be further investigated in future studies. iii) While RPA shows sensitivity to changes in consciousness, its clinical utility should be benchmarked against established metrics like the Bispectral index (BIS) and Patient State Index (PSI) from the monitoring apparatuses used in the clinical settings. iv) Expanding the application of RPA to diverse ADHD subtypes, comorbidity symptoms, as well as varying age and sex groups, is critical for generalizing its utility as a biomarker and understanding the heterogeneity of the disorder. Additionally, finding a more rigorous correlation between various phenotype data and RPA results will help identify what behavioral characteristics are influenced by the differences in RPA. v) Our general-coupled oscillator model framework has many potentials to be refined, for example, by utilizing more biologically informative models such as Wilson-Cowan model to reveal more information of the biophysical mechanism and the mode transition dynamics. Additionally, the observation that relative phase patterns and robust statistical features emerge consistently across multiple frequency bands, particularly in the beta and theta ranges, suggests the need to expand our current narrow-band model into a more comprehensive wide-band framework. vi) Lastly, in David Marr’s levels of analysis where a model consist of computational, algorithmic, and implementational levels [66, 67] – each level corresponding to questions about what problems are addressed, how the problem is addressed algorithmically, and how the mechanism is biophysically implemented – our model is at the level of implementation. Only the expansion and combinations of our model with the algorithmic and computational levels will provide a complete picture of how the information is processed via the top-down and bottom-up switching in our brain with respect to the external world and internal model construction.

### Conclusion

This study introduces RPA as a powerful and versatile tool for investigating the brain dynamics, offering new insights into brain state transitions, hierarchical processing, and neuropsychiatric conditions. Our findings demonstrate that the fluctuating interplay between top-down and bottom-up processing is an intrinsic property of large-scale brain dynamics, reinforcing a dynamical systems perspective in which the brain continuously reconfigures its information processing organization. Our study offers a new research avenue exploring how external stimuli, cognitive demands, and pathological conditions influence this delicate balance. By integrating RPA with experimental neuroscience, we can investigate how spontaneous directional fluctuations shape perception, cognition, and consciousness, ultimately advancing our understanding of the fundamental principles of neural computation.

## 4 Methods

### 4.1 Relative Phase Analysis of EEG signals

In electroencephalography (EEG), a participant wears a cap that has tens or hundreds of electrodes attached to it. These electrodes record the brain’s electrical activity over time. A key challenge in neuroscience is to understand how the brain processes external and internal information. To address this problem, we try to investigate information flow from the electrical activity in the brain using the relative phase dynamics approach.

EEG signals can be decomposed into amplitude and phase components over time using the Hilbert transform [68]. The phase of each electrode reflects the specific timing of its signal within a periodic waveform. To determine the directionality of information flow, we use the relative phase, which is derived by subtracting the global mean phase from the phase of each electrode [5, 7]. In the following equations, *θ*_*j*_ represents the signal phase of *j*^th^ electrode, and Θ denotes the system’s mean phase.

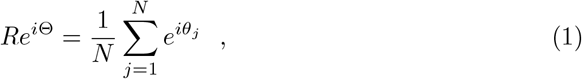

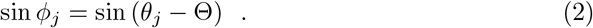

Here, Eq. (1) represents the global mean phase and Eq. (2) represents the relative phase. Eq. (1) defines the system’s phase denoted as Θ, which represents the global mean phase of the system at each time point, capturing the brain’s overall phase state. In Eq. (2), we subtract the system’s phase from each signal’s phase, and define it as relative phase *ϕ*_*j*_. When the relative phase is a positive value, it means that the signal from an electrode leads the global mean phase, whereas a negative value indicates that it lags the global mean phase. To increase the accuracy of the relative phase estimation, we take the sine of *ϕ*_*j*_. This transformation allows us to obtain continuous relative phase values ranging from *−*1 to 1, utilizing only the values from the imaginary axis [22]. Using this approach, we can reliably estimate whether *ϕ*_*j*_ is relatively phase-leading or lagging at each time point.

### 4.2 *K*-means clustering of the relative phase Patterns

The relative phase in the brain shows distinct patterns over time, reflecting different brain states. We found that the dynamics of the relative phase patterns show different aspects depending on the brain states. Interestingly, these patterns are not random in conscious states but follow specific dynamics. The *K*-means clustering algorithm finds dominant patterns as centroids by iteration in a given dataset [69, 70]. We performed *K*-means clustering to find dominant patterns in the relative phase dynamics.

*K*-means clustering divides the given data into *K* clusters, a parameter that must be predetermined. To determine the optimal *K* for resting state brain patterns, we evaluate the clustering performance using the Davies-Bouldin Index (DBI) [71]. The DBI reflects clustering quality, with higher values when the average distance between the same cluster is greater than the average distance between different clusters, otherwise lower values. After examining the DBI values across varying *K* values, we found that the optimal number of clusters for the brain’s resting state is *K* = 4 (see Figure S5 in the Supplementary Information).

In the main text, we analyzed differences between different groups or brain states. To investigate these differences, we identified universal relative phase patterns based on resting state data. We derived four centroids using *K*-means clustering from two datasets: the concatenated resting state recordings of 18 subjects from the University of Michigan (UofM), and the concatenated resting state recordings of the Healthy Brain Network (HBN) control group (66 subjects), respectively. We excluded the NKI dataset due to noise introduced by simultaneous fMRI-EEG recordings. Despite analyzing different datasets, we found that the same four centroids emerged consistently: i) front-leading/back-lagging pattern (mode 1), ii) left-leading/right-lagging pattern (mode 2), iii) right-leading/left-lagging pattern (mode 3), iv) back-leading/front-lagging pattern (mode 4) (see Figure S6 in the Supplementary Information). Therefore, we analyzed relative phase dynamics using these four centroids as standard.

### 4.3 Multiple linear regression using the universal centroids

The *K*-means clustering algorithm assigns every time frame to one of the four modes, even if some ambiguous frames do not closely resemble any mode. This limitation introduces ambiguity, as frames that fall between distinct patterns are still forced into one cluster. To address this issue, we performed multiple linear regression with the universal centroids [72]. Unlike *K*-means, which provides binary assignments, multiple linear regression assigns continuous weights to each mode for every time frame, representing the contribution of each pattern more accurately.

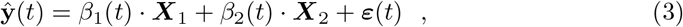

where **ŷ** is the relative phase pattern at a given time *t*, ***X***_1_ and ***X***_2_ are regressors based on the universal centroids that indicate front-leading/back-lagging (averaged with mode 1 and reverse mode 4) and left-leading/right-lagging patterns (averaged with mode 2 and reverse mode 3), respectively (see Supplementary Information Section B.3 for detail). *β*_1_ and *β*_2_ represent the coefficients of explanation using each regressor. *ε* represents residuals that cannot be explained by the superposition of ***X***_1_ and ***X***_2_. As a result, the coefficients are given for each time frame.

Quantifying brain states is challenging because both the relative phase time series and traditional measurements are calculated for each channel. Conventional approaches often reduce the dimension of the time series by averaging the measurements across channels, even though this approach may overlook spatially specific brain activity. In contrast, our approach preserves spatial information by focusing on temporal patterns without averaging over channels. In summary, we can obtain three distinct coefficient time series (i.e. *β*_1_, *β*_2_, and |***ε***|) using multiple linear regression with the universal centroids.

### 4.4 Statistics over brain states

By applying multiple linear regression and comparing whether |*β*_1_| is larger than |*β*_2_|, we can obtain an index for each time frame. This index indicates the similarity to specific modes; i) mode 1 (front-leading/back-lagging, i.e. positive sign of *β*_1_), ii) mode 2 (left-leading/right-lagging, i.e. positive sign of *β*_2_), iii) mode 3 (right-leading/left-lagging, i.e. negative sign of *β*_2_), iv) mode 4 (back-leading/front-lagging, i.e. negative sign of *β*_1_). Using these indices, we can construct a probability density function to calculate the mode distribution over different brain states. In addition, by weighting each time frame with its corresponding regression coefficient, we can better distinguish the brain states (see Supplimentary Information Section B.7). To get the information intuitively comparing with unweighted version, we normalize the statistics with the mean of 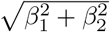 in baseline (eyes-closed resting state) to reasonably compare across different brain states in Figures S18 and S20. This weighted approach enhances the effects of clear patterns and reduces the effects of ambiguous patterns. In the following analyses of EEG relative phase patterns, we find that the mode distribution can distinguish different brain states.

Furthermore, we can examine the dwell time for each mode. The dwell time refers to the time spent in a mode before transitioning to another. The mode time series changes noisily at the boundary of each mode. By investigating the average dwell time distribution across each mode, we can reduce the impact of these boundary effects and obtain a more accurate characterization of brain states. In addition, we can also define the weighted dwell time by multiplying the coefficients of multiple linear regression.

For the general anesthesia dataset, where sample sizes varied significantly across states (resting = 90, LOC = 15, Burst = 30, Suppression = 30, ROC = 45), we applied bootstrap resampling 1,000 times per state and compared the resampled means. To compare mode occurrences, dwell times, and iPR across different states, we performed one-way ANOVA for each mode. For each resampling iteration, confidence intervals were computed, and *p*-values were estimated as the proportion of bootstrap samples in which the confidence interval did not contain zero. To account for multiple comparisons, *p*-values were adjusted using the Bonferroni-Holm method. For the typically developed vs. ADHD dataset, where the sample sizes were sufficiently large and the difference less pronounced (typically developed = 66, ADHD = 40), we conducted statistical analyses without resampling. We performed a two-sample t-test to compare mode occurrences and dwell times. We refer to the Supplementary Information Section A for details on sample sizes.

### 4.5 Principal Component Analysis (PCA) of relative phase dynamics

Principal component analysis (PCA) is a linear dimensionality reduction technique that finds the dominant axes of variation from a given dataset [73, 74]. By applying PCA, we can obtain the principal components, with each component explaining a proportion of the total variance along its corresponding axis. The inverse Participation Ratio (iPR) quantifies the heterogeneity of a given distribution by measuring how many principal components significantly contribute to the overall variance [75, 76]. We can use the iPR to find an effective number of the principal axes across different brain states and quantify the differences by comparing them.

The relationship between linear regression, *K*-means clustering, and PCA has been previously studied and indicated [77]: It was demonstrated that the cluster centroid space of the continuous solution of the *K*-means clustering algorithm spans the first *K* − 1 principal directions. We applied all three methods to check the consistency of our results. This provides robust validation by cross-checking the findings obtained through each technique.

### 4.6 EEG Data & preprocessing

In this section, we describe the dataset and preprocessing pipeline (For details, see the dataset description in Supplementary Information A). We used three types of datasets: 1) general anesthesia EEG data (UofM, University of Michigan) to find dominant relative phase patterns in the sub-second time scale because anesthesia is an obvious stimulus to weaken consciousness; 2) checkerboard task of simultaneous EEG-fMRI data (NKI, Nathan Kline Institute) to examine the meaning of the relative phase patterns as compared with functional networks based on fMRI BOLD signals; 3) ADHD/Control EEG data (HBN, Healthy Brain Network) to suggest that the relative phase approach can be applied in detecting neurodevelopmental disorders. For the preprocessing procedure, we used the EEGLAB package in MATLAB [78]. The EEG signal was processed as follows: 1) notch filtered at 60 Hz to remove power line noise, 2) bad channels were detected and removed using the trimOutlier.m function, and 3) band-pass filtered across six frequency bands (delta: 1-4 Hz, theta: 4-8 Hz, alpha: 8-12 Hz, low beta: 12-20 Hz, high beta: 20-30 Hz, gamma: 30-50 Hz).

### 4.7 fMRI Data & preprocessing

In fMRI BOLD signals, we use the preprocessed data served by the reference site [25, 79, 80] (func pp filter gsr sm0.mni152.3mm.nii.gz). Briefly, the structural preprocessing included skull-stripping, tissue segmentation (gray matter, white matter, cerebrospinal fluid), and registration to the MNI152 template. The functional preprocessing included slice timing, motion correction, boundary-based co-registration, nuisance regression (Friston’s 24 head motion parameters, white matter, cerebrospinal fluid, and global signal), band-pass filter (0.01-0.1 Hz) and linear and quadratic detrending (For the more details on preprocessing see Supplementary Information Section A). For spatial averaging under the 1,000-parcel Schaefer parcellation scheme, fMRI volumes were resampled to 2 mm isotropic resolution to align with the parcellation template. Voxel-level time series were then averaged within each parcel, and the resulting parcel-wise time series were Z-scored across time.

### 4.8 Other measures

To validate that the relative phase approach is analogous and related to other measures, we compared it with key metrics such as amplitude and phase-lag measures. Additionally, we compared relative phase with measures that quantify traveling wave propagation to demonstrate that our measure captures the directionality of activity. To accomplish this, we applied the relative phase and other measures to the EEG data collected from the University of Michigan and looked at how the relationships between measures change across different states.

For each state, we segmented the alpha-band filtered signal into non-overlapping epochs of 2 seconds (see Figures S2, S3, and S4 for results of 0.5, 5, and 10 second time windows). The measures were then computed over these data segments. Finally, pairwise linear correlations were calculated for the time series of the measures. These correlation values were Fisher Z-transformed and averaged across subjects. P-values were calculated from these values and corrected for multiple comparisons using the Bonferroni method.

#### 4.8.1 Amplitude

To calculate the amplitude, we applied the Hilbert transform to the segmented signals and averaged the instantaneous amplitude over each window. The amplitude time series for each channel was then averaged across the frontal and parietal regions. For the frontal region, we selected 14 channels corresponding to the electrodes Fpz, Fp1-2, AF3-4, AF7-8, Fz, F1-4, and F7-8. Similarly, for the parietal region, we considered the electrodes Pz, P1-8, POz, PO3-4, and PO7-8.

#### 4.8.2 Phase Lag Index

As a measure of functional connectivity, we used the weighted phase-lag index (wPLI), which takes into account the strength of the phase difference between signals and is robust to volume conduction and uncorrelated noise sources [22]. The wPLI ranges from 0 (random phase lead/lag) to 1 (consistent phase lead/lag). After obtaining the wPLI matrix for all channel pairs, we averaged the values for the frontal, parietal, and fronto-parietal regions. Additionally, we defined a directional connectivity measure using the signed wPLI, averaging over the frontal and parietal channels.

#### 4.8.3 Traveling wave propagation

We calculated two measures of traveling waves using three lines of electrodes spanning from the frontal to the occipital regions. These included one line of midline electrodes and two lines of electrodes positioned to the left and right of the midline, respectively. First, we quantified traveling wave propagation using 2D fast Fourier transform (FFT) representations of the time-by-channel data matrix [10]. The spectral peaks in the first and fourth quadrants of the 2D FFT spectra correspond to forward- or backward-propagating waves. We took the maximum value, *P*_real_, over spatial frequencies in the temporal frequency domain, and repeated this procedure with shuffled electrodes to obtain a surrogate measure as *P*_shuffled_. The amount of waves was then calculated in decibels [dB] as 10 · log_10_ (*P*_real_*/P*_shuffled_), which represents waves relative to the null distribution. The forward- or backward-propagating waves were calculated for each electrode line, and then averaged. Additionally, we measured traveling wave propagation by calculating the phase gradient over the three lines of electrodes [8]. This involved computing the relative phase across the selected electrodes and fitting a polynomial surface to these values. For each time point, we extracted the y coefficient from the polynomial, which quantifies the phase gradient from the frontal to the occipital electrodes.

### 4.9 Simulation of relative phase dynamics with Kuramoto model

To elucidate the mechanism behind the mode fluctuations, we ran simulations of relative phase dynamics using a Kuramoto model, a well-known canonical coupled oscillator model frequently used to model large-scale cortical dynamics [7, 24, 81–94]. Using the generated relative phase signals from the model on human cortical network, we constructed the spatially extrapolated relative phase topographic map to obtain regression coefficients (*β*_1_ and *β*_2_) that specify the mode distribution over different brain states, using the same procedure for the empirical EEG data. Specifically, we focused on modeling the general anesthesia EEG data (UofM, University of Michigan) to find the model parameters that best match the empirical statistics of the brain states including mode occurrence and dwell time. Below, we describe the model and simulation procedure in more detail.

#### 4.9.1 Structural connectivity data

The structural and function connectome data was collected from 70 healthy individuals on a 3T scanner [95]. We used a group-averaged structural network generated with a consensus approach [96] which preserve the edge length distribution in individual participants, described in detail elsewhere [46, 97, 98]. In particular, we used the network with 114 regions. The structural matrix was further binarized, yielding a binary adjacency matrix of size 114 by 114 specifying the connectivity between brain regions. This matrix was then used to specify the connection between nodes in the Kuramoto model, explained below.

#### 4.9.2 Kuramoto model

To simulate the phase signals of individual regions on the brain network, we utilized a Kuramoto model with a generalized coupling function that includes a constant term (*c*_0_), which results in a more realistic first order approximation of many coupling functions obtained from the phase reduction method [27, 28, 57]. The model is written as:

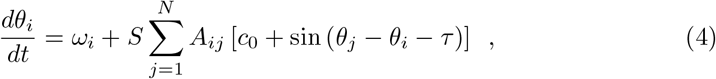

where *θ*_*i*_ is the phase of the oscillator *i, ω*_*i*_ is the intrinsic frequency of the oscillator, *S* is the global coupling strength, **A** is the binary adjacency matrix of the network, and *τ* is the phase shift term between two nodes capturing the transmission delay between regions. *c*_0_ is a constant that results from the first-order approximation of the coupling function. Seen in another way, this term can be thought of as a natural consequence of incorporating diffusive coupling in limit cycle oscillators [29, 44, 45, 59]. For all of our simulations, we set *ω* = 10 Hz for all nodes to focus on the alpha band frequency, and *τ* = 0.12*π*, which approximately translates into the average time delay between brain regions.

#### 4.9.3 Ornstein-Uhlenbeck (OU) and other stochastic processes in *c*_**0**_

The previous study has shown that inclusion of the *c*_0_ parameter in the coupling function of oscillators results in a rich repertoire of phase dynamics, especially relevant to the transitions between two distinct and dominant modes in the brain: one in which high-degree midline cortical regions phase-lag the network becoming sink of the directionality, and one in which they phase-lead the network and becoming source of the directionality [45]. To simulate the signals, we assumed that the brain exhibits random fluctuations in the *c*_0_ parameter. The stochastic dynamics were specified with the Ornstein-Uhlenbeck (OU) process [99], a commonly adopted framework for implementing time-varying random noise [100–104]. Formally, the OU process for our variable of interest *x*_*t*_ (= *c*_0_ at time *t*) is described as:

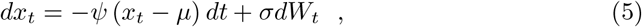

where *ψ* is the parameter controlling for the rate of drift toward the constant *µ* which specifies the mean of the process, and *σ* is the parameter controlling the variance of the Gaussian noise *dW*_*t*_. While *µ* could be a free parameter, we set *µ* = sin (*τ*) for all simulations as this corresponds to the in-phase synchronous state.

To explore other potential random dynamics in *c*_0_ that might account for the observed relative phase dynamics, we tested two additional stochastic processes involving pitchfork bifurcations. More specifically, the *supercritical* case was described by its normal form involving third-order term, with the variable centered at *µ* as follows:

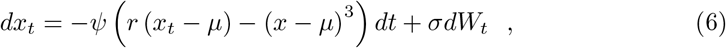

where *r >* 0 is the parameter controlling the distance between the two stable fixed points (as shown in Supplementary Figure S26B). Note that the two stable fixed points appear only when *r* > 0 [105].

For the *subcritical* case, an additional fifth-order term was included to stabilize the dynamics as follows:

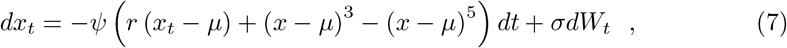

where *r* is a parameter bounded between −0.25 *< r <* 0 to allow for two stable fixed points (Supplementary Fig. S26C). For all simulations, we chose *r* = 0.1 for the supercritical case and *r* = *−*0.18 for the subcritical case to allow for two stable fixed points on both sides of *µ* = sin (*c*_0_), namely, the attractors states for mode 1 and 4 (Supplementary Figure S25A).

#### 4.9.4 Parameter estimation using grid search

To estimate the parameters of the stochastic processes, we ran simulations of Kuramoto oscillators with stochastic processes in *c*_0_ for 60s with a time step of Δ*t* = 0.001 (for a total of 60,000 time steps). This was repeated for 10 unique random seeds, and the resulting mode occurrences and dwell time were averaged. We used grid search to run simulations with the given values of *ψ, σ* (step size of 0.5), and the coupling strength *S* (step size of 0.1), and selected the set of parameters which exhibited the minimum mean absolute error from the empirical mode occurrences and dwell times.

## Data availability

The EEG dataset for the general anesthesia presented in this article are not publicly available. Requests to access these datasets should be directed to J.-Y.M. The simultaneous EEG-fMRI dataset with the checkerboard task, and ADHD vs. typically developed EEG dataset are publicly available: for more details, please refer to the references [25, 40].

## Code availability

Code for models, simulations and analyses is available at https://github.com/youngjai/RelativePhaseAnalysis [106].

## Acknowledgments

This work was supported by IBS-R015-Y3 (to J.-Y.M., Y.C., Y.K.) from the Institute for Basic Science of Korea (IBS), by the Basic Science Research Program through the National Research Foundation of Korea (NRF) funded by the Ministry of Education of Korea (RS-2023-00272652; to Y.P.), the National Research Foundation of Korea (NRF) grant funded by the Korea Government (MSIT) (NRF-2022R1C1C1007095, RS-2023-00217361, RS-2024-00398768; to S.-J.H.), by the DeepTech TIPS funded by the Ministry of SMEs and Startups (RS-2023-00302377; to H.K.), by the National Institutes of Mental Health of USA (R01-MH119099 to C.J.H. and R01-2P50MH109429 subawarded to C.J.H.). We thank Haesung Lee for his help in the numerical simulations. We thank Dongho Kim for his advice on the framework of the analysis. We thank Heonsoo Lee for his advice in analyzing the general anesthesia dataset. We thank Qawi K. Telesford for his advice in preprocessing and analyzing simultaneous EEG-fMRI dataset. We thank Jong-eun Lee for his advice in analyzing the ADHD dataset. We thank Eunhee Ji for her advice and comments on fMRI analysis and results. We thank Sunhyun Min, Seoyeon Kim, Jehyeop Lee, Minseo Cho, Jiwon Kim, and F. D. Poohy for their support in carrying out the study.

## Author contributions

Y.P., Y.C., H.K., J.-Y.M. designed the study, and Y.K., J.H.W., H.C., T.X., U.L., S.-J.H., C.J.H. provided input and revisions. Y.P., H.K., J.-Y.M. developed the pipeline of the relative phase analysis. Y.P., Y.C., H.K., Y.K., J.-Y.M. developed and implemented the data analysis. J.H.W., J.-Y.M. developed and conducted the numerical simulations. Y.C., Y.K., H.C. performed the statistical analysis. G.A.M., T.X., U.L., S.-J.H. provided datasets. Y.P., Y.C., Y.K., J.H.W., J.-Y.M. wrote the manuscript, which was revised by H.K., H.C., U.L., S.-J.H., C.J.H.

## Additional information

### Supplementary information

This manuscript is accompanied by supplementary material, containing additional texts and figures.

**Correspondence and requests for materials** should be addressed to Joon-Young Moon.

