## Supplementary Information for "Sub-Second Fluctuation between Top-Down and Bottom-Up Modes Distinguishes Diverse Human Brain States"

**C Model Analysis: Additional Kuramoto Model Analyses** **22**

### A Metheods: Data Collection & Preprocessing Procedure

#### A.1 General anesthesia EEG dataset

EEG recordings from eighteen healthy volunteers (20-40 years old) were collected at the University of Michigan [1, 2]. The study was reviewed in accordance with the recommendations of the Institutional Review Boards specializing in human subject research at the University of Michigan, Ann Arbor (Protocol #HUM0071578). Written informed consent was obtained from all participants in accordance with the Declaration of Helsinki. Nine participants underwent general anesthesia, the others were recorded without anesthesia. EEG data were recorded with 128-channel HydroCel nets, Net Amps 400 amplifiers (Electrical Geodesic, Inc., USA) at a sampling rate of 500 Hz.

The experimental paradigm was designed to simulate the conditions of general anesthesia typically used in a surgical setting. Participants in the anesthesia group initially received propofol through increasing infusion rates over three consecutive 5-minute blocks: **Block 1 (100 µg/kg/min)**, **Block 2 (200 µg/kg/min)**, and **Block 3 (300 µg/kg/min)**. Responsiveness was assessed every 30 seconds using the verbal command, **"Squeeze your left/right hand twice,"** with left/right assignment randomized. Following this, isoflurane was administered with air and 40% oxygen at **1.3 age-adjusted minimum alveolar concentration (MAC)**. Isoflurane was maintained for **3 hours** before being discontinued.

In the EEG data, we extracted 5-minute eyes-open and 5-minute *eyes-closed* resting states from the no anesthesia group (nine subjects). In the general anesthesia group (nine subjects), we extracted five types of brain states over time to study differences among the different brain states; 1) 5-minute *eyes-closed* resting state before general anesthesia, 2) 3-minute *LOC* (loss of consciousness) state after LOC marker, 3) *burst* state pieces with a total of 5-minute, 4) *suppression* state pieces with a total of 5-minute, and 5) 5-minute *ROC* (recovery of consciousness) state after ROC marker. LOC and ROC markers in the EEG data were measured by responding to the verbal command ("Squeeze your left/right hand twice" with left/right randomized) every 30 seconds [1]. In the data, six subjects of the anesthesia group showed burst and suppression states under anesthesia. The burst state is a period when the power in the spectrogram suddenly increases across all frequency bands even though the subjects are under anesthesia. On the contrary, in the suppression state, the power remains extremely low. In particular, we manually extracted the burst and suppression states because it is unclear to quantitatively define the states. To increase sample size for state comparisons, we divided the data into 1-minute segments and analyzed them.

#### A.2 Data preprocessing in the UofM (University of Michigan) dataset

Before the main analysis, we performed the following preprocessing procedure using the EEGLAB v2022.1 package in MATLAB [3]: i) To remove 60 Hz as power line noise, we perform notch filtering with the range 59 to 61 Hz using the `pop_eegfiltnew.m` function. ii) To also remove the global trend in a low-frequency band and artifact noise in a high-frequency band, we perform detrending with the range 0.5 to 100 Hz using the `pop_eegfiltnew.m` function. iii) In the EEG, some electrodes have anomalous signals due to the bad connection. For more accurate analysis, we remove bad channels for each state of each subject. We detect bad channels across all states of all subjects using the `trimOutlier.m` function with the range 0.0001 to 100 μV. Then, we define a common set of bad channels identified in common across all states of all subjects. After excluding the common bad channel set, we detect and remove bad channels for each state of each subject using the `trimOutlier.m` function with the same range. Additionally, if the number of the removed bad channels in a state of a subject is greater than 25% of the total number of channels, we exclude the state from the main analysis. In the main text, we analyze 18 *eyes-closed*, 5 *LOC*, 6 *burst*, 6 *suppression*, and 9 *ROC* states. iv) Finally, we perform six types of bandpass filters (delta: 1-4 Hz, theta: 4-8 Hz, alpha: 8-12 Hz, low beta: 12-20 Hz, high beta: 20-30 Hz, gamma: 30-50 Hz) using the `pop_eegfilternew.m` function.

#### A.3 NKI (Nathan Kline Institute) simultaneous EEG-fMRI dataset

Over the past several decades, many researchers have discovered that different brain regions are activated depending on the task. The activation of the brain regions requires a higher blood oxygen level because the firing of neurons requires more energy. For these reasons, BOLD (blood-oxygen-level-dependent) is commonly used as an indirect measurement when we investigate brain analysis. In contrast, the meaning of EEG dynamics patterns is less clear than fMRI because the EEG is measured on the surface of the scalp. However, fMRI data has worse temporal resolution than EEG. Because EEG and fMRI are complementary to each other, studying simultaneous EEG-fMRI recordings has recently begun to attract attention. In this study, we compare EEG relative phase patterns with fMRI BOLD signals to discover the meaning of the EEG relative phase patterns.

The dataset is collected at the NKI (Nathan Kline Institute) [4]. The recordings were obtained from 22 individuals between the ages of 23 and 51 years old. EEG data contains 64-channels using a customized Brain Products (BrainCapMR consisting of 61 cortical channels, two EOG channels placed below (channel 63) and above (channel 64) the left eye, and one ECG channel (channel 32)). All individuals were consented in accordance and compliance with the Institutional Review Board (IRB) at NKI. fMRI data was acquired using a 12-channel head coil on 3T Siemens TIM Trio. All BOLD fMRI sequences were acquired with these following parameters: TR=2100ms; TE=24.6ms; slices=38; matrix size=64×64; voxel size=$3.469 \times 3.469 \times 3.330 \text{mm}^3$.

#### A.4 Data preprocessing in NKI simultaneous EEG-fMRI dataset

To analyze the simultaneous EEG-fMRI dataset, we perform several steps of preprocessing procedure in both EEG and fMRI data, respectively. Basically, we download the dataset in [4, 5]. In EEG signals, we preprocess the data using EEGLAB: 1)

Due to the RF signal of MR scanner, we performed gradient artifact removal using `pop_fmrib_fastr.m`. 2) We removed the pulse artifact because the MR scanner strengthens the heartbeat signal. (QRS/heartbeat detection using the ECG channel and `pop_fmrib_qrsdetect.m`; Pulse artifact removal using `pop_fmrib_pas.m`). 3) We conduct bandpass filter with alpha band (8-12 Hz) using `pop_firws.m` in EEGLAB. In fMRI BOLD signals, we use the preprocessed data served by the reference site [5, 6] (`func_pp_filter_gsr_sm0.mni152.3mm.nii.gz`). Briefly, the structural preprocessing included skull-stripping, tissue segmentation (gray matter, white matter, cerebrospinal fluid), and registration to the MNI152 template. The functional preprocessing included slice timing, motion correction, boundary-based co-registration, nuisance regression (Friston's 24 head motion parameters, white matter, cerebrospinal fluid, and global signal), band-pass filter (0.01-0.1 Hz) and linear and quadric detrending.

#### A.5 ADHD inattentive subtype and control dataset

To analyze the differences in the directionality of phase relationship between the ADHD group and the control group, we used eyes closed and eyes open resting state data from the Healthy Brain Network (HBN) dataset released by the Child Mind Institute [7]. The HBN dataset is a large-scale data collection targeting adolescents (aged 5-21) living in the New York City area. It consists of extensive multimodal datasets covering a wide spectrum of commonly encountered clinical psychopathologies including attention deficit hyperactivity disorder (ADHD). To examine the real-time directionality of phase relationship across whole brain regions, we used high-density electroencephalography (EEG) data recorded at a sampling rate of 500 Hz using a 128-channel geodesic hydrocel system by Electrical Geodesics Inc. (EGI).

To establish a more robust sample with ADHD-Inattentive type, we selected individuals who had been diagnosed with ADHD inattentive type without comorbidity. The control group consists of individuals who had not received any diagnosis. The EEG signal in the alpha spectrum peak shows a qualitative difference between age under and above 10 [8]. Therefore, we chose individuals aged 11 and older. In conclusion, resting state data from a total of 53 ADHD participants (41 male, mean age 14.11, standard error (s.e.) 0.27) and 88 control individuals (44 male, mean age 14.07, s.e. 0.28) were used for the analysis. Participants are encouraged to discontinue stimulant medication due to its potential effects on cognitive and behavioral testing and brain function mapping [7]. However, those who choose not to or are required by their physicians to continue medication can still participate, with their medication intake recorded on the day of the study. Three participants in the selected ADHD-inattentive group were on medication during data collection, and they were not excluded from the analysis.

#### A.6 Data preprocessing in HBN dataset

For a more accurate analysis, we applied a 60 Hz noise reduction to resting state data using a notch filter through the MATLAB function `eeglab_notchFilter.m` in the EEGLAB toolbox. After noise reduction, participants with data recording issues or excessively high noise levels were excluded from the analysis. Subsequently, we used

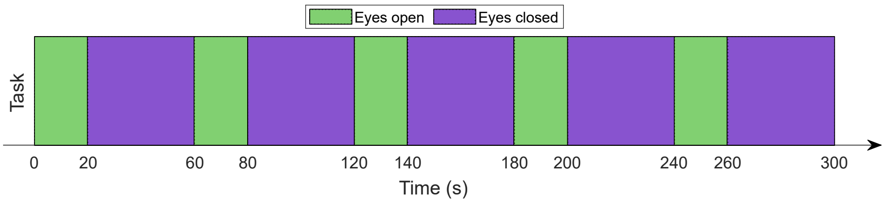

**Fig. S1 Task description of HBN. Five cycles of alternating conditions.** 20 seconds of eyes open periods, followed by 40 seconds of eyes closed periods.

a band-pass filter with a cutoff frequency range of 0.5 Hz to 100 Hz through the MATLAB function `pop_eegfiltnew.m`. We then identified low-quality electrodes using MATLAB function `trimOutlier.m` with a lower bound of 0.0001 standard deviation and an upper bound of 100 standard deviations. After identifying bad channels across all subjects from both the control group and the ADHD group, any channel that was deemed a bad channel in 25% or more of the total subjects was designated as a common bad channel. Considering the bad channel results and the symmetry of channel locations, we eliminated 8 channels (8, 14, 21, 25, 48, 67, 77, 119) that were common across all subjects. After excluding the common bad channels, we reused the `trimOutlier.m` function to identify and remove additional bad channels for each subject, applying the same criteria. Subjects with fewer than 93 remaining channels were excluded from the analysis.

After noise reduction and bad channel removal, the final analysis comprised 40 participants in the ADHD group (34 males, mean age 13.97, s.e. 0.29) and 66 participants in the Control group (34 males, mean age 14.06, s.e. 0.32). Using the same function `pop_eegfiltnew.m`, we extracted the alpha spectrum (8 Hz to 12 Hz) and then separated the eyes open and eyes closed resting states. The EEG resting state data in the HBN dataset were collected over a 5-minute duration, involving five cycles of alternating conditions: 20 seconds of eyes open followed by 40 seconds of eyes closed (see Figure S1). To minimize the influence of the preceding state, we analyzed only specific segments of the data. We extracted 16.8 seconds for the eyes open state (2.2 seconds at the beginning and 1 second at the end). For the eyes closed state, we extracted 33.6 seconds (5.2 seconds at the beginning and 1 second at the end) for our analysis.

### B Data Analyses: Additional Relative Phase Analyses

#### B.1 Other measures of different scales

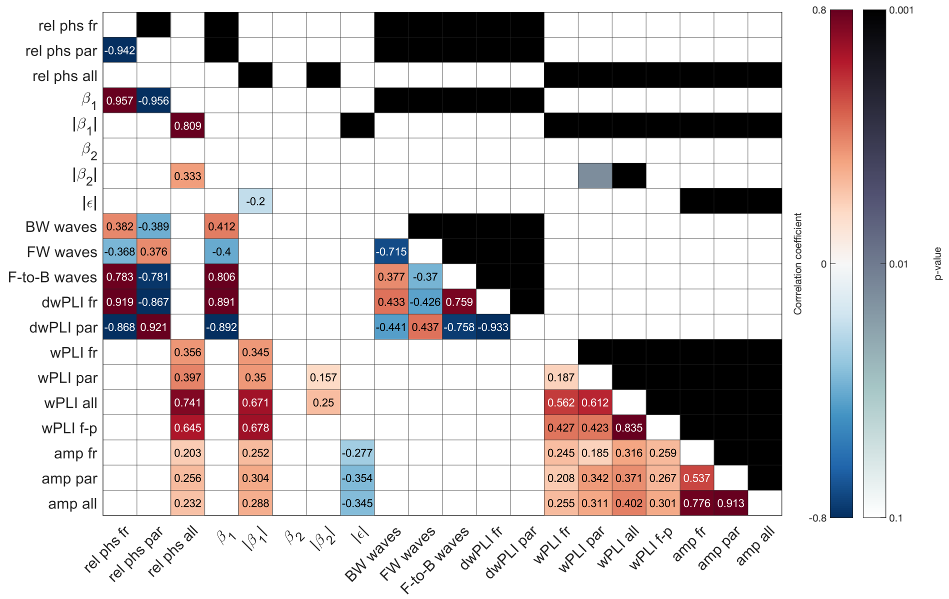

**Fig. S2 Correlation matrices for 0.5-second time windows in the eyes-closed state, with correlation values on the lower triangle and $p$-values on the upper triangle.** The relative phase averaged over frontal regions shows a positive correlation of 0.919 with directional wPLI over frontal regions ($p < 0.001$), and a correlation of 0.801 with frontal-to-posterior (F-P) traveling waves ($p < 0.001$). Similarly, the relative phase averaged over parietal regions shows a positive correlation of 0.921 with directional wPLI over parietal regions ($p < 0.001$), and a negative correlation of $-0.803$ with frontal-to-posterior traveling waves ($p < 0.001$).

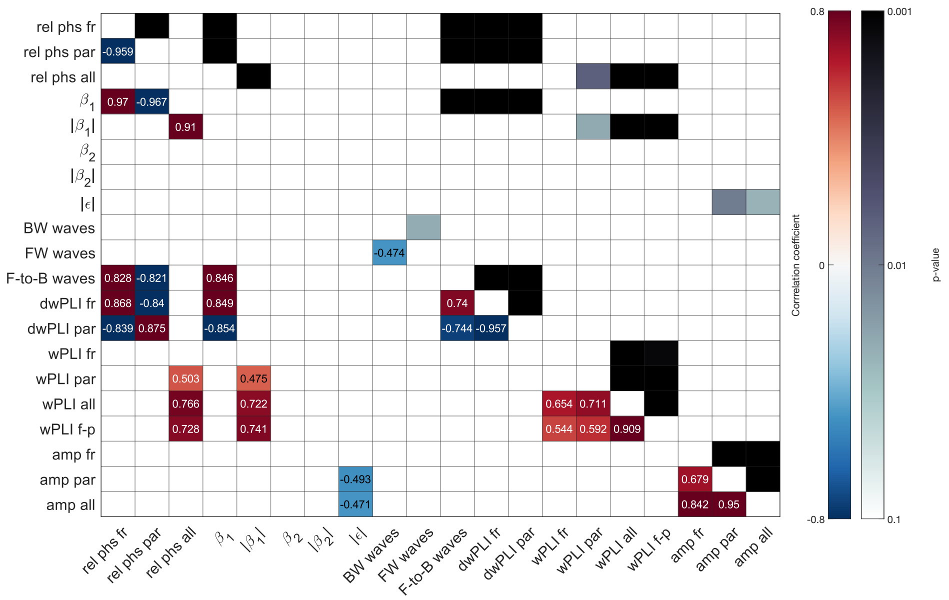

**Fig. S3 Correlation matrices for 5-second time windows in the eyes-closed state, with correlation values on the lower triangle and $p$-values on the upper triangle.** The relative phase averaged over frontal regions shows a positive correlation of 0.868 with directional wPLI over frontal regions ($p < 0.001$), and a correlation of 0.828 with frontal-to-posterior (F-P) traveling waves ($p < 0.001$). Similarly, the relative phase averaged over parietal regions shows a positive correlation of 0.875 with directional wPLI over parietal regions ($p < 0.001$), and a negative correlation of $-0.821$ with frontal-to-posterior traveling waves ($p < 0.001$).

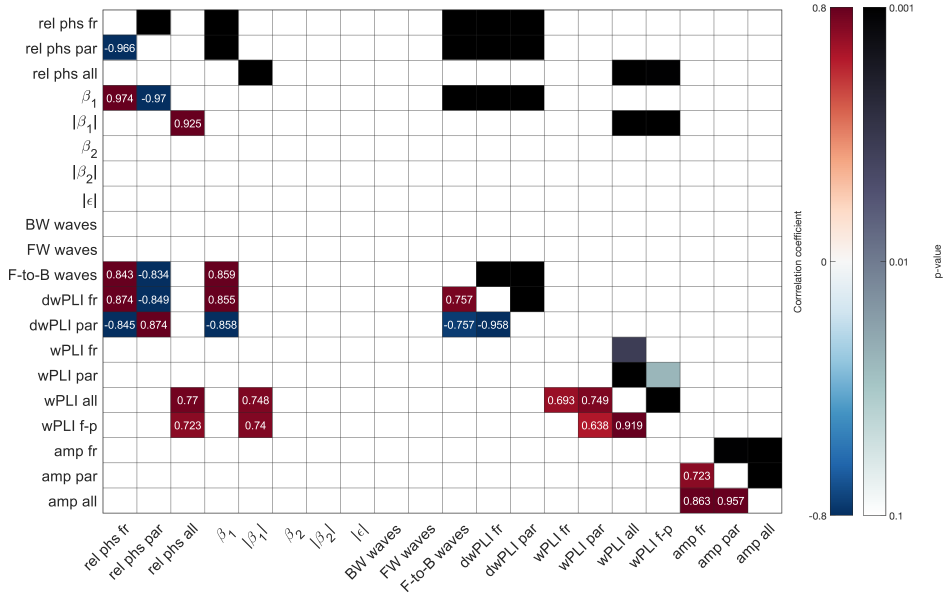

**Fig. S4 Correlation matrices for 10-second time windows in the eyes-closed state, with correlation values on the lower triangle and $p$-values on the upper triangle.** The relative phase averaged over frontal regions shows a positive correlation of 0.874 with directional wPLI over frontal regions ($p < 0.001$), and a correlation of 0.861 with frontal-to-posterior (F-P) traveling waves ($p < 0.001$). Similarly, the relative phase averaged over parietal regions shows a positive correlation of 0.874 with directional wPLI over parietal regions (p¡0.001), and a negative correlation of $-0.857$ with frontal-to-posterior traveling waves ($p < 0.001$).

#### B.2 Optimal number of $K$-means clustering algorithm in UofM dataset

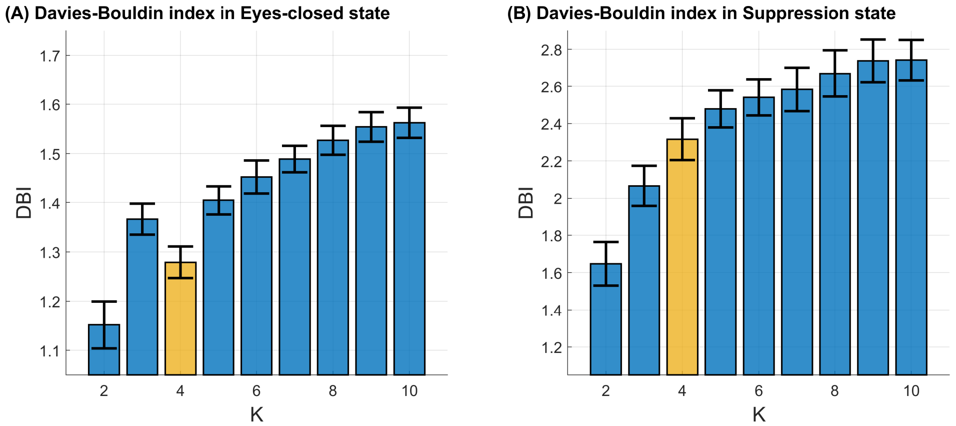

**Fig. S5 Davies-Bouldin index as increasing $K$ in the general anesthesia data.** To determine the optimal $K$ in the human brain EEG recordings, we calculate the Davies-Bouldin index in a baseline state as (A) *eyes-closed* state (because of the resting state) and (B) *suppression* state. The bar indicates the mean value of the eighteen subjects' indexes and the error bar is the standard error. As a result, we set the optimal $K$ as 4 (orange bar) because we find the global optimum at $K = 4$ in the resting state.

#### B.3 Universal centroids from UofM and HBN datasets

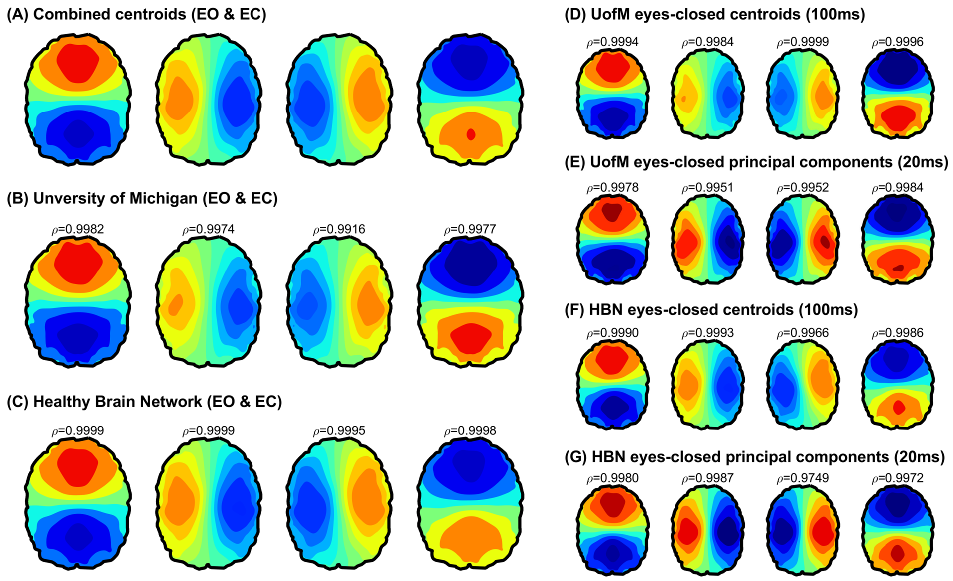

**Fig. S6 Topographic maps of universal centroids using UofM and HBN datasets.** In each dataset, there are two types of resting states (eyes-open and eyes-closed); (A-C) the averaged centroids of both eyes open and eyes-closed resting states in each dataset. $\rho$ in (B-C) indicates the correlation coefficient with combined centroids. (D-G) the centroids of 100ms and 20ms time resolution and $\rho$ indicate the correlation coefficient between two different time resolutions. All centroids show almost the same. Since the differences in centroids across datasets are minimal, we use the combined centroids in the main analysis for consistency.

#### B.4 Optimal time scale in the EEG relative phase time series in the UofM dataset

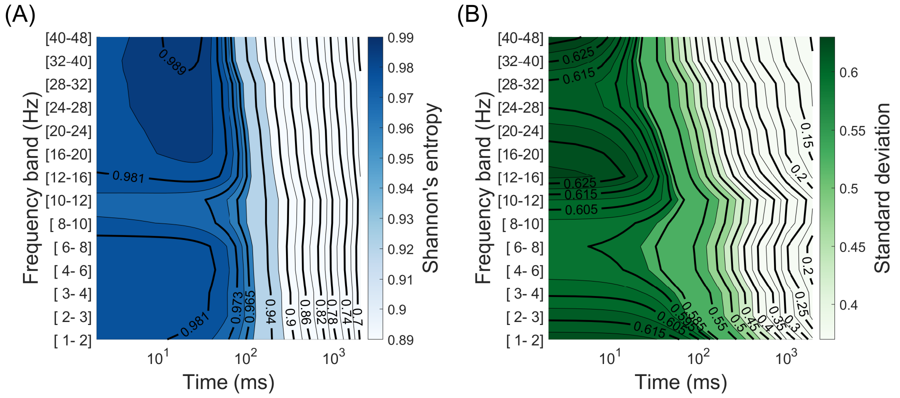

**Fig. S7 Optimal time scale of the relative phase time series in the UofM dataset.** To reduce data size without the loss of information, we investigate the channel-wise averaged (A) Shannon's entropy and (B) standard deviation of downsampled time series in the eyes-closed resting state according to frequency bands. In particular, we find that the information is preserved at a larger time scale in the alpha band (8-12 Hz). As a result, we analyze the relative phase dynamics with downsampled as 100 ms in the main text.

#### B.5 Mode distribution with considering ambiguous frame as "Other"

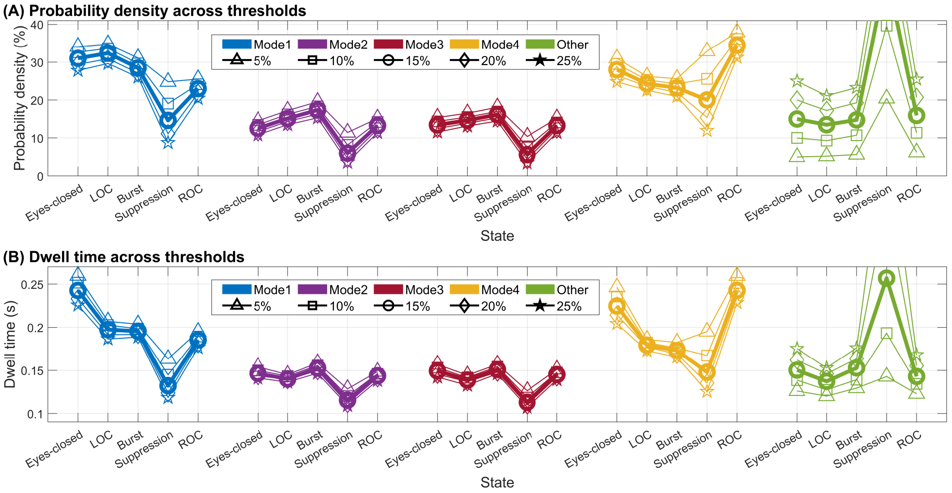

**Fig. S8 Mode occurrence and dwell time considering "Other".** Probability density (A) and dwell time (B) across different brain states, considering various "Other" threshold criteria. To clarify the relationship between relative phase dynamics and the level of consciousness, we classify ambiguous frames as "Other" based on the residual, $|\varepsilon|(t)$.

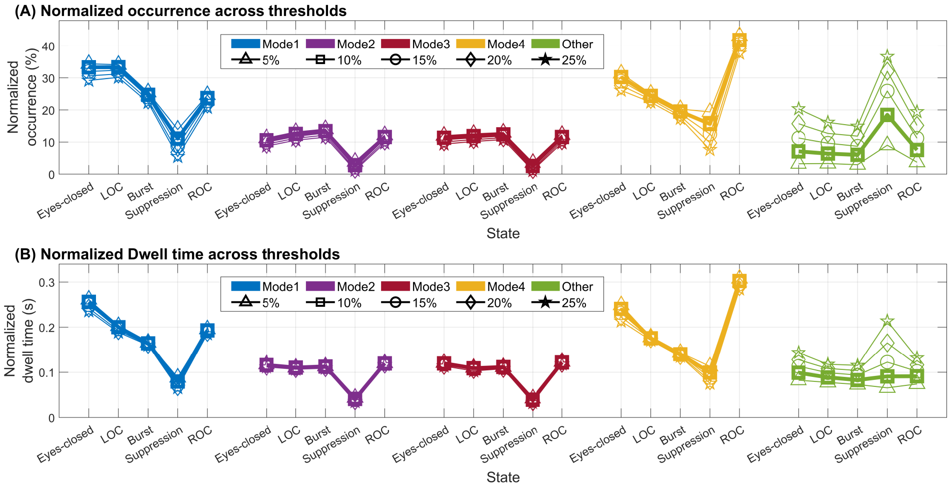

**Fig. S9 Normalized mode occurrence and dwell time considering "Other".** Normalized occurrence (A) and normalized dwell time (B) across different brain states, considering various "Other" threshold criteria. To clarify the relationship between relative phase dynamics and the level of consciousness, we classify ambiguous frames as "Other" based on the residual, $|\varepsilon|(t)$.

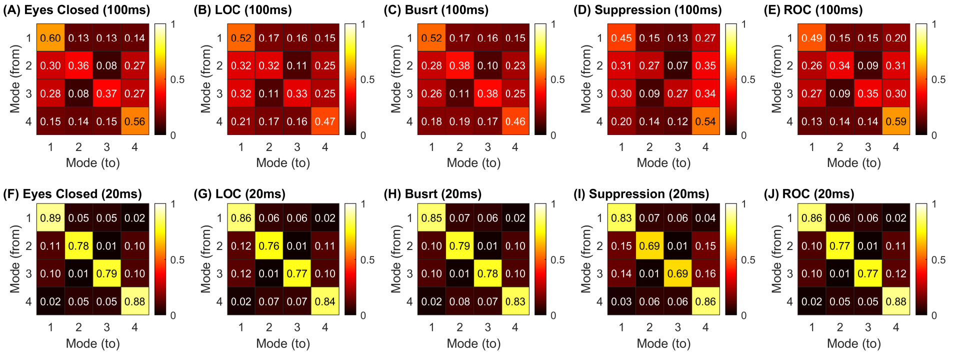

**Fig. S10 Transition matrices across different anesthetic states.** As the level of consciousness decreases (e.g. the suppression state), the transitions between modes become more frequent, indicating reduced stability within a single mode.

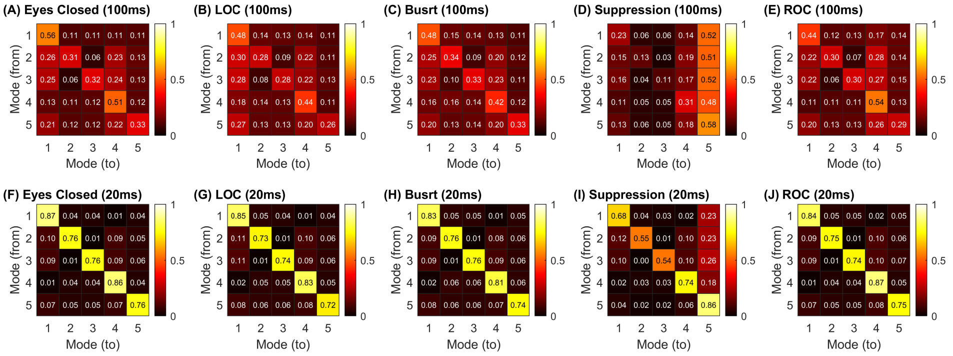

**Fig. S11 Transition matrices across different anesthetic states, with introducing the "Other" state.** As the level of consciousness decreases (e.g. the suppression state), the brain mode spends more time in "Other" state, which means that the brain wave patterns become random and fail to form stable patterns.

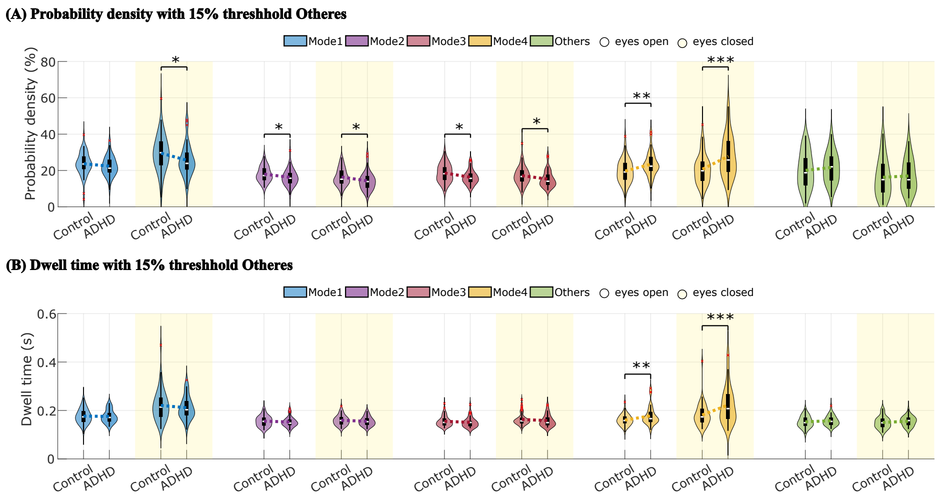

**Fig. S12 HBN data's probability density (A) and dwell time (B) of the HBN data with a 15% threshold for "Others," based on the eyes-closed state of the control group.** The results are generally similar to those in Figures 4B and 4C. In both (A) and (B), a significant difference between the ADHD group and the control group is observed in the eyes-closed state, specifically in mode 4, at the $p < 0.001$ level. For the remaining cases: (A): Two-sample t-test, t-value ($p$-value) = 2.07 (0.041), 2.03 (0.045), 2.16 (0.033), 2.51 (0.014), 1.99 (0.050), −2.97 (0.004), −3.82 (0.0002), (B): Two-sample t-test, t-value ($p$-value = −2.99 (0.004), −3.52 (0.0007).

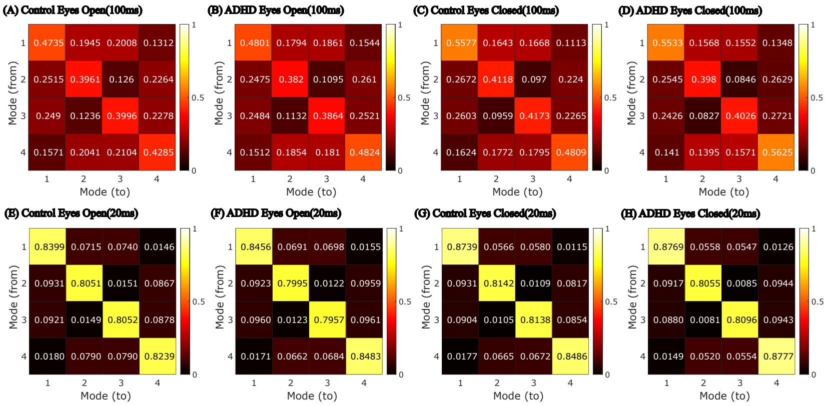

**Fig. S13 Transition matrices of the Control group and the ADHD group in the resting state.** The values of the matrices show differences between groups in both 100 ms and 20 ms time windows. As seen in the main figure, the largest difference between the two groups occurs in the transition from mode 4 to mode 4, with a more significant disparity observed in the 100 ms window compared to the 20 ms window: mode 4 to mode 4 two-sample t-test results, t-value ($p$-value) (1) eyes open, 100 ms $-3.28$ ($p < 0.01$), (2) eyes closed, 100 ms $-3.64$ ($p < 0.001$), (3) eyes open, 20 ms $-2.96$ ($p < 0.01$), (4) eyes closed, 20 ms $-3.10$ ($p < 0.01$). Additionally, the transition matrices indicate that transitions between mode 1 and mode 4 often pass through mode 2 and mode 3. Similarly, mode 2 and mode 3 do not transition directly between each other. This tendency is more clearly evident in the 20 ms transition matrix.

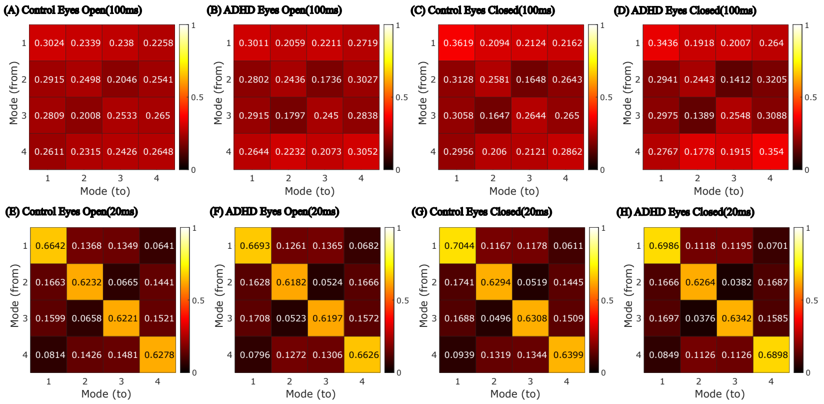

**Fig. S14 Transition matrices of the weighted calculated version for the Control group and the ADHD group in the resting state.** The values of the matrices show that the differences between the two groups have generally decreased compared to the unweighted version. In the 100 ms window, for both eyes-open and eyes-closed states, the transition from mode 4 to mode 4 still shows a significant difference between the two groups at the $p < 0.001$ level (t-value, eyes-open $-3.33$, eyes-closed $-4.01$). However, in the 20 ms window, the differences between the two groups are significant at the $p < 0.01$ level for both states (t-value, eyes-open $-3.38$, eyes-closed $-3.99$). Likewise, in the weighted version, the transition matrices show that transitions between mode 1 and mode 4 frequently pass through mode 2 and mode 3. This pattern is more pronounced in the 20 ms transition matrix.

#### B.6 Mode properties in different frequency bands

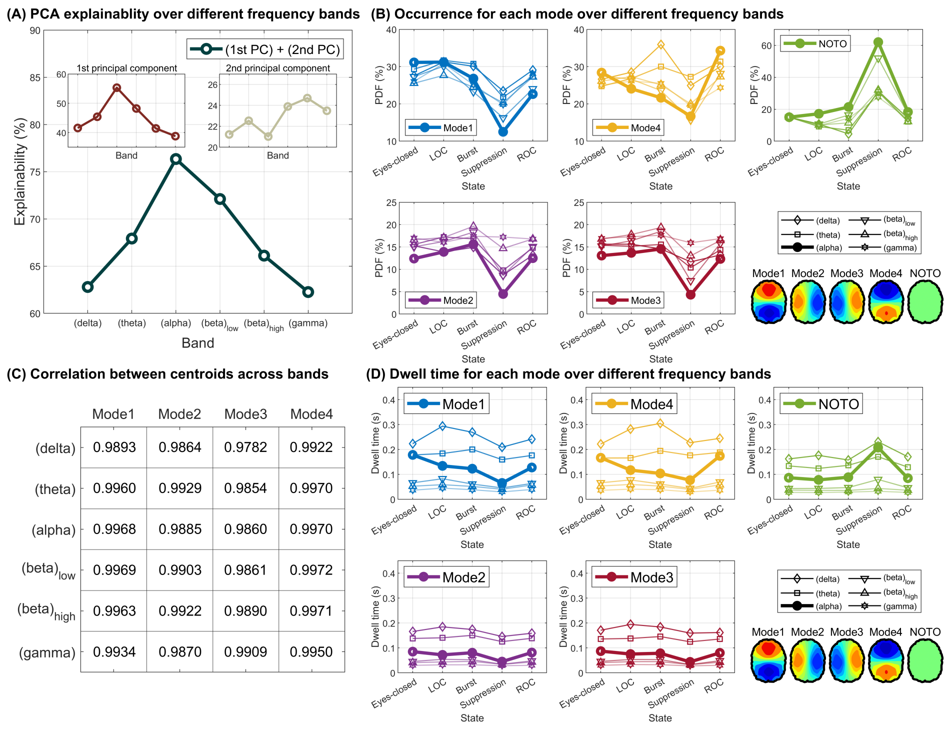

| | Mode1 | Mode2 | Mode3 | Mode4 |
|---|---|---|---|---|
| (delta) | 0.9893 | 0.9864 | 0.9782 | 0.9922 |
| (theta) | 0.9960 | 0.9929 | 0.9854 | 0.9970 |
| (alpha) | 0.9968 | 0.9885 | 0.9860 | 0.9970 |
| $(beta)_{low}$ | 0.9969 | 0.9903 | 0.9861 | 0.9972 |
| $(beta)_{high}$ | 0.9963 | 0.9922 | 0.9890 | 0.9971 |
| (gamma) | 0.9934 | 0.9870 | 0.9909 | 0.9950 |

**Fig. S15 Relative Phase Analysis across different frequency bands.** (A) The cumulative explainability of the first two principal components reaches its maximum in the alpha (8-12 Hz) band, suggesting that our methodology is most effective in the alpha band. (B) The mode occurrence trend across different levels of consciousness is also prominent in the alpha band. (C) Despite variations in frequency range, we find that the dominant patterns remain consistent across each band. (D) The dwell time trend follows the mode occurrence pattern.

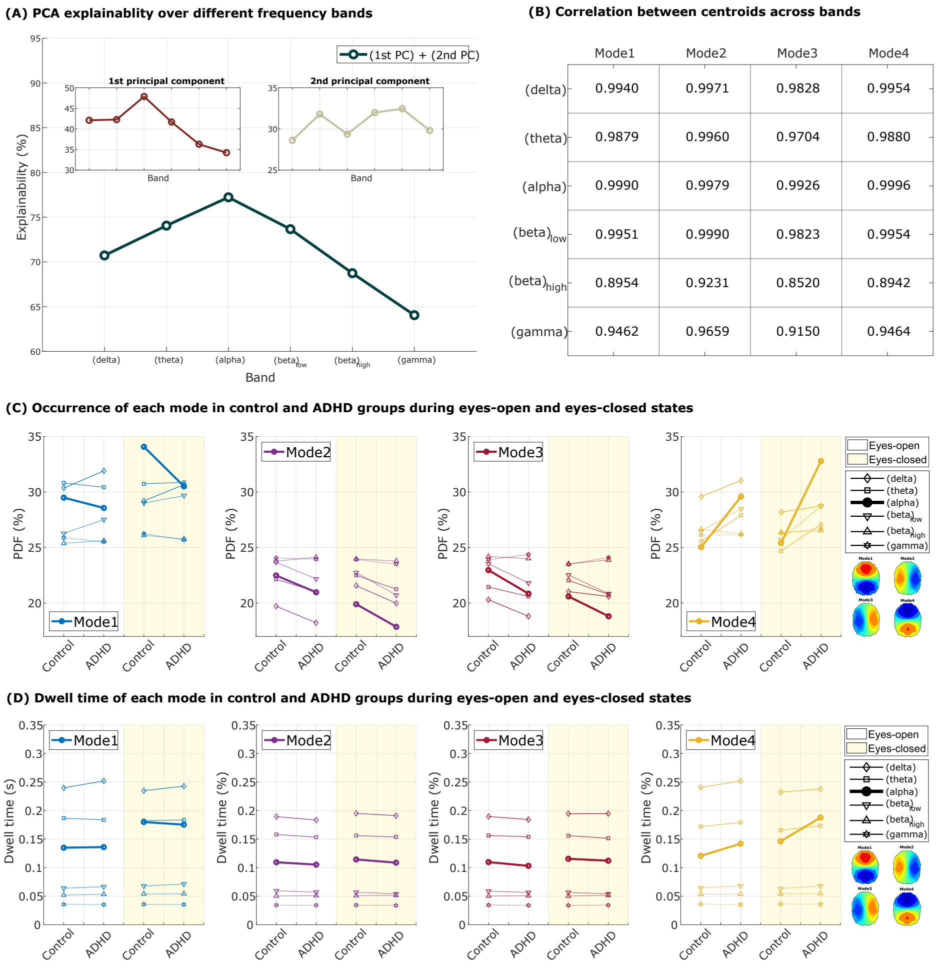

**Fig. S16 Relative Phase Analysis across different frequency bands for the control and ADHD group.** (A) The cumulative explainability of the first two principal components reaches its maximum in the alpha (8-12 Hz) band, similarly demonstrating that the information value is highest in the alpha band. (B) Although the frequency ranges vary, the dominant patterns remain stable across all bands. (C) The mode occurrence trend of the control and ADHD groups for both eyes-open and eyes-closed states. The pattern of a higher mode ratio in mode 4 for the ADHD group is also observed in other bands. Two sample t-test results t-value ($p$-value): **Mode 1**: $-2.46$ (0.02, theta, EO), $-2.54$ (0.01, theta, EC), 2.09 (0.04, alpha, EC), **Mode 2**: 2.09 (0.04, low beta, EC), **Mode 3**: 2.09 (0.04, alpha, EO), **Mode 4**: $-3.55$ ($< 0.001$, alpha, EO), $-3.75$ ($< 0.001$, alpha, EC), $-2.60$ (0.01, low beta, EC), (D) Although weaker than in the PDF, the same pattern in mode 4 is also observed in dwell time. two sample t-test results t-value ($p$-value) **Mode 1**: $-2.16$ (0.03, theta, EC), **Mode 4**: $-3.56$ ($< 0.001$, alpha, EO), $-4.09$ ($< 0.001$, alpha, EC), $-2.09$ (0.04, low beta, EO), $-2.32$ (0.02, low beta, EC).

#### B.7 100 ms/20 ms time-scale results and unweighted/weighted results for Relative Phase Analysis of alpha band

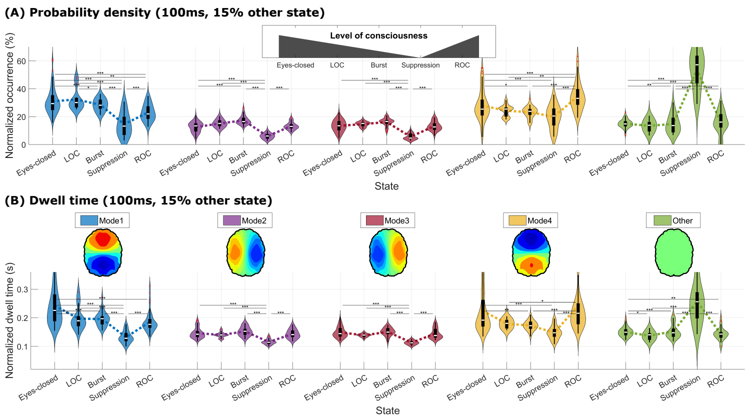

**Fig. S17 Relative Phase Analysis for 100 ms time resolution and unweighted mode properties.** (A) The probability density function varies across different levels of consciousness. In particular, modes 1 and 4 are related to the depth of consciousness, suggesting that the occurrence of these modes may serve as key markers for monitoring brain states. (B) The dwell time, which represents the duration spent in each mode, also differs across brain states, showing distinct trends depending on the level of consciousness.

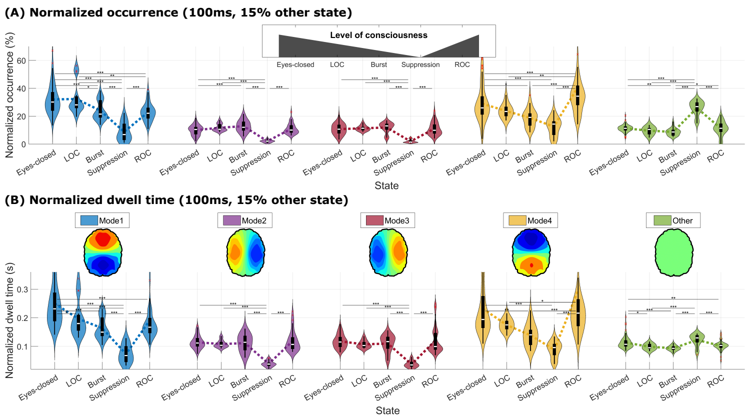

**Fig. S18 Relative Phase Analysis for 100 ms time resolution and weighted mode properties.** (A) The probability density function varies across different levels of consciousness. In particular, modes 1 and 4 are related to the depth of consciousness, suggesting that the occurrence of these modes may serve as key markers for monitoring brain states. (B) The dwell time, which represents the duration spent in each mode, also differs across brain states, showing distinct trends depending on the level of consciousness. We normalize the statistics using the mean of $\sqrt{\beta_1^2 + \beta_2^2}$ in baseline (the *eyes-closed* resting state) to provide a meaningful unit, allowing for reasonable comparison across different brain states.

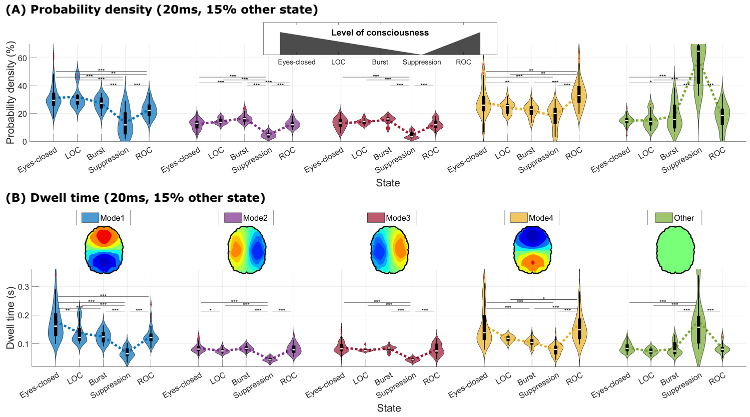

**Fig. S19 Relative Phase Analysis for 20 ms time resolution and unweighted mode properties.** (A) The probability density function varies across different levels of consciousness. In particular, modes 1 and 4 are related to the depth of consciousness, suggesting that the occurrence of these modes may serve as key markers for monitoring brain states. (B) The dwell time, which represents the duration spent in each mode, also differs across brain states, showing distinct trends depending on the level of consciousness.

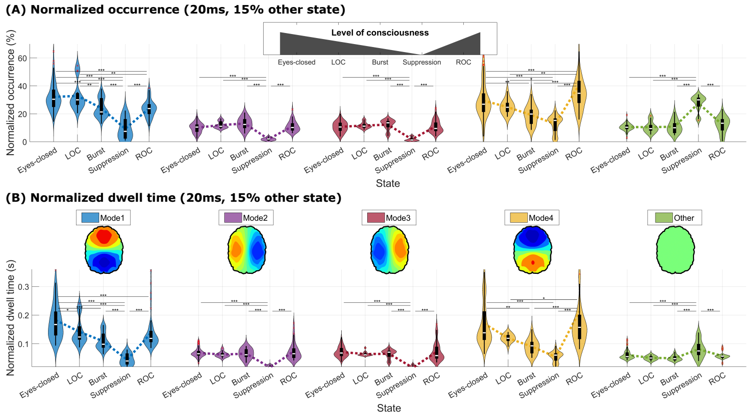

**Fig. S20 Relative Phase Analysis for 20 ms time resolution and weighted mode properties.** (A) The probability density function varies across different levels of consciousness. In particular, modes 1 and 4 are related to the depth of consciousness, suggesting that the occurrence of these modes may serve as key markers for monitoring brain states. (B) The dwell time, which represents the duration spent in each mode, also differs across brain states, showing distinct trends depending on the level of consciousness. We normalize the statistics using the mean of $\sqrt{\beta_1^2 + \beta_2^2}$ in baseline (the *eyes-closed* resting state) to provide a meaningful unit, allowing for reasonable comparison across different brain states.

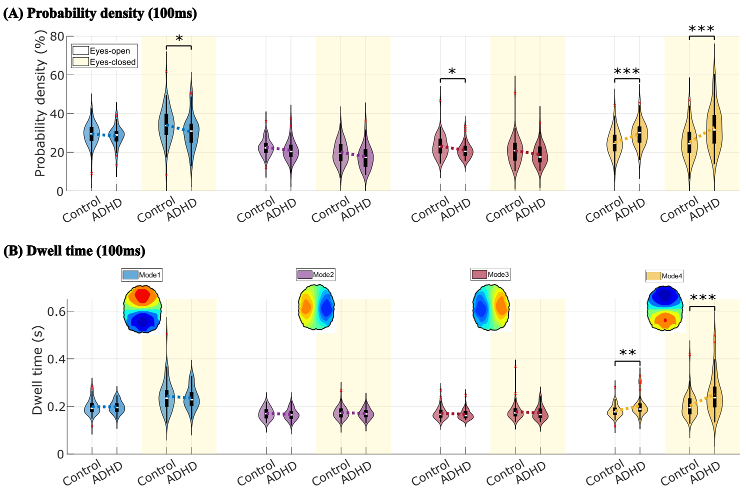

**Fig. S21 Relative Phase Analysis for a 100 ms time window, alpha, and unweighted mode properties in the control and ADHD groups.** (A) Probability density figure, a violin plot version of Figure 4B. Two sample t-test results t-value ($p$-value): **Mode 1**: 2.12 (0.04, EC), **Mode 3**: 2.14 (0.04, EO), **Mode 4**: −3.67 (< 0.001, EO), −4.19 (< 0.001, EC). (B) Dwell time figure, a violin plot version of Figure 4C. Two sample t-test results t-value ($p$-value): **Mode 4**: −3.23 (0.002, EO), −3.89 (< 0.001, EC).

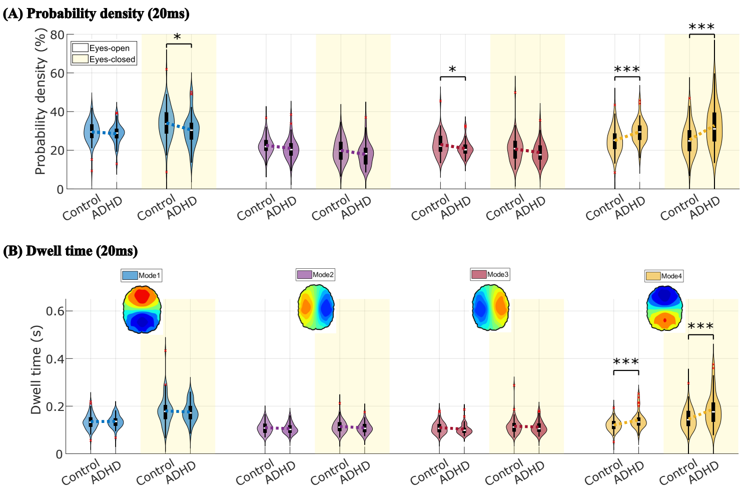

**Fig. S22 Relative Phase Analysis for a 20 ms time window, alpha, and unweighted mode properties in the control and ADHD groups.** (A) Probability density figure. Two sample t-test results t-value ($p$-value): **Mode 1**: 2.09 (0.04, EC), **Mode 3**: 2.09 (0.04, EO), **Mode 4**: −3.56 (< 0.001, EO), −4.09 (< 0.001, EC). (B) Dwell time figure. Two sample t-test results t-value ($p$-value): **Mode 4**: −3.55 (< 0.001, EO), −3.75 (< 0.001, EC).

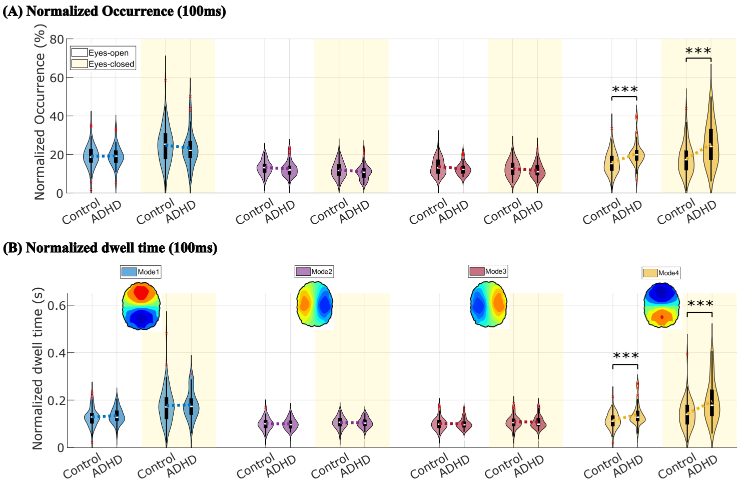

**Fig. S23 Relative Phase Analysis for a 100 ms time window, alpha, and weighted mode properties in the control and ADHD groups.** (A) Probability density figure. Two sample t-test results t-value ($p$-value): **Mode 4**: $-3.82$ ($< 0.001$, EO), $-4.18$ ($< 0.001$, EC). (B) Dwell time figure. Two sample t-test results t-value ($p$-value): **Mode 4**: $-3.40$ ($< 0.001$, EO), $-3.83$ ($< 0.001$, EC).

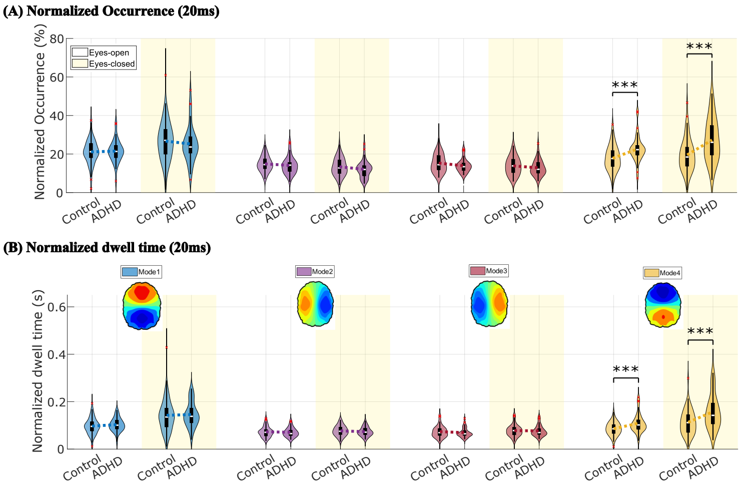

**Fig. S24 Relative Phase Analysis for a 20 ms time window, alpha, and weighted mode properties in the control and ADHD groups.** (A) Probability density figure. Two sample t-test results t-value ($p$-value): **Mode 4**: $-3.65$ ($< 0.001$, EO), $-4.05$ ($< 0.001$, EC). (B) Dwell time figure. Two sample t-test results t-value ($p$-value): **Mode 4**: $-3.57$ ($< 0.001$, EO), $-3.69$ ($< 0.001$, EC).

### C Model Analysis: Additional Kuramoto Model Analyses

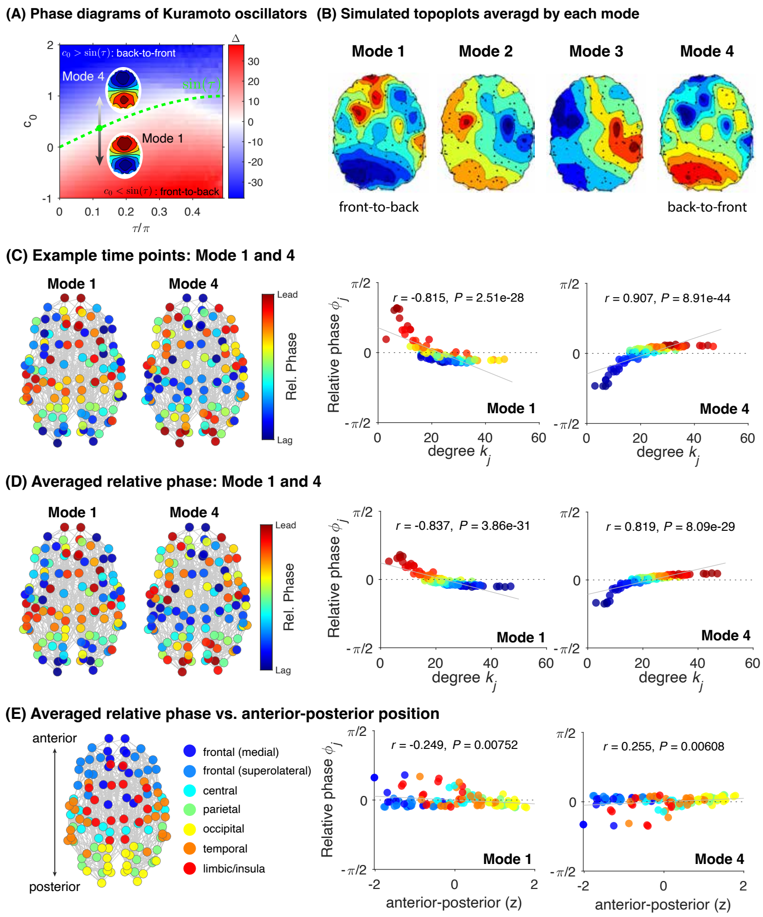

**Fig. S25 Simulated Relative phase dynamics in Kuramoto coupled oscillators with various types of attractor dynamics.** (A) Phase diagram of $\Delta$ $(= \omega - \Omega)$ in the coupled oscillator system, shown in colors, indicating the difference between the intrinsic frequency ($\omega$) and the frequency of the population during the steady state ($\Omega$). Diagrams are plotted as a function of phase shift term $\tau$ and the $c_0$ parameter. $\Delta > 0$ (red region) indicates parameter space where, for the non-drifting, partially locked or synchronous oscillators, higher-degree regions phase-lag (containing Mode 1). $\Delta < 0$ (blue region) indicates parameter space where higher-degree regions phase-lead (containing Mode 4). Green lines indicate $\sin(\tau)$, corresponding to the in-phase synchronous state. (B) Simulated topographic maps by each mode, averaged across all simulated instances. (C) Visualization of relative phase on the brain network (left) and scatter plots (right) of relative phase as a function of node degree $k_j$ during example time points for modes 1 and 4 (shown in Figure 5B). Reported are Pearson's correlation coefficients and p-values. Colors indicate relative phase of each node (red: phase-lead; blue: phase-lag). (D) Same plots as in (C) but for relative phases averaged across all time points corresponding to mode 1 or mode 4. (E) Relationship between the anterior-posterior coordinate (z-scored) and the relative phase of each node. Colors indicate corresponding brain areas.

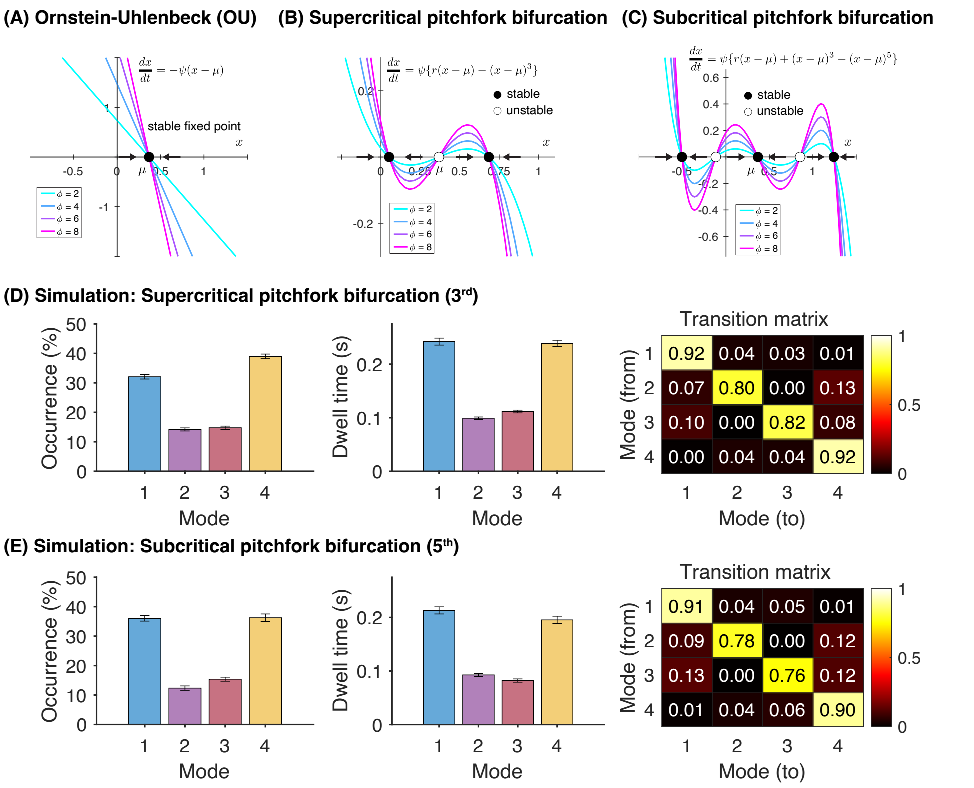

**Fig. S26 Stochastic processes with additional attractors capture the front-to-back and back-to-front relative phase dynamics observed in the empirical data.** (A-C) Vector fields for three different types of attractor dynamics. Ornstein-Uhlenbeck (OU) process (A), shown in Figure 5, has a stable fixed point at $\mu$. Supercritical pitchfork bifurcation with third-order term (B), and subcritical pitchfork bifurcation with third- or fifth-order terms (C) have additional stable fixed points around $\mu$. (D) Statistics of empirical mode distributions, simulated by the random process in $c_0$ with supercritical pitchfork bifurcation (third-order) illustrated in (B). Conventions are the same as those shown in Figures 5C and D (UofM general anesthesia dataset). Plotted are the distribution of occurrences (left), dwell time (middle), and transition matrices among the four brain states (right). Simulation parameters: $r = 0.1$, $\psi = 11$, $\sigma = 1$, $S = 1.8$. (E) Same plots as in (D) but showing simulated results using the subcritical pitchfork bifurcation (fifth-order) illustrated in (C). Simulation parameters: $r = -0.18$, $\psi = 10$, $\sigma = 11$, $S = 0.9$.
